# Un*lox*ing the assembly and activation mechanism of Cre recombinase using Cryo-EM

**DOI:** 10.1101/2021.07.31.454597

**Authors:** Kye Stachowski, Andrew Norris, Devante Potter, Vicki Wysocki, Mark P. Foster

**Affiliations:** Department of Chemistry and Biochemistry, The Ohio State University, Columbus, Ohio, USA 43210; Resource for Native Mass Spectrometry Guided Structural Biology, The Ohio State University, Columbus, Ohio, USA 43210

**Author notes:** Corresponding Author: Mark P. Foster).

**Keywords:** Cre recombinase, Cryo-EM, DNA bending, native mass spectrometry

## Abstract

Mechanistic understanding of the structural basis for DNA recombination in the Cre-*loxP* system has largely been guided by crystallographic structures of tetrameric synaptic complexes (intasomes). These structural and biochemical studies have suggested that conformational changes and DNA bending in presynaptic complexes underlie site-selection and activation mechanisms of Cre recombinase. Here we used protein engineering and various DNA substrates to isolate the Cre-*loxP* (54 kDa), Cre2-*loxP* (110 kDa), and Cre4-*loxP*2 assembly intermediates, and determined their structures using cryo-EM to resolutions of 3.9 Å, 4.5 Å, and 3.2 Å, respectively. Progressive DNA bending along the assembly pathway enables formation of increasingly intimate protein-protein interfaces. Insufficient stabilization of important protein motifs observed during the assembly process provides a compelling explanation for the observed half-the-sites activity, and preferential bottom strand cleavage of *loxP* sequences. We found that selection of *loxP* sites is largely dependent on Cre’s ability to bend and stabilize the spacer region between two recombinase binding elements. Application of 3D variability analysis to the tetramer data reveals a propensity for motion along the pathway between protomer activation and Holliday junction isomerization. These findings help us to better understand *loxP* site specificity, controlled activation of alternating protomers, the basis for the observed bias of strand cleavage order, and the importance of conformational sampling, especially with regards to site-selection and activity among Cre variants. Furthermore, our findings provide invaluable information for the rational development of designer, site-specific recombinases for use as gene editing technologies.

**Highlights:**

- Cryo-EM structures of Cre-*loxP* assembly intermediates (monomer, dimer, and tetramer) provide insights into mechanisms of site recognition, half-the-sites activity, strand cleavage order, and concerted strand cleavage
- Selectivity of *loxP* sites arises from few base-specific contacts made by each protomer and is mainly driven by formation of phosphate contacts and DNA deformations that are maximal in the fully assembled “active” tetramer
- *Cis* and *trans* interactions of the β2-3 loop (i) define which sites are “active” and (ii) ensure half-the-sites activity
- Protein flexibility plays a central role in enabling DNA sequence scanning, recruitment of a second protein to form a dimer, synapsis, control of activity, as well as subsequent recombination steps
- Conformational sampling within the tetrameric complex was uncovered using 3D variability analysis and revealed the importance of protein-protein interfaces for site-selection and activation of Cre-*loxP* complexes

## Introduction

We report Cryo-EM structural studies of DNA recognition and assembly of recombination intermediates by the enzyme Cre (<u>c</u>auses <u>re</u>combination). Cre, a tyrosine site-specific DNA recombinase (YSSR), enables precise removal or replacement of defective DNA through a highly controlled process that precludes DNA legions^1, 2^, elevating its candidacy as an effective gene editing technology.^3–7^ Cre excises, exchanges, or inverts double stranded DNA beginning with recognition of two 34-bp asymmetric *loxP* DNA sites (Figure 1). Each *loxP* site comprises two palindromic 13-base pair (bp) recombinase binding elements (RBEs) that are recognized in antiparallel fashion by two Cre proteins. The RBEs are separated by an 8-bp asymmetric spacer that determines the orientation of each site, and the outcome of a recombination reaction.^8–11^ Four molecules of Cre assemble with two *loxP* DNA sites and, in alternating fashion, tyrosine residues of each protomer perform a series of strand cleavages, exchanges and ligations, forming covalent 3’-phosphotyrosine and Holliday junction intermediates in order to generate recombinant products. Cre catalyzes this reaction without the need of additional factors or consumption of ATP, and thus, has emerged as a powerful tool for genome engineering in the laboratory.^12^ However, expanded use of the technology, in particular for targeting arbitrary or asymmetric recognition sites is hindered by our limited understanding of the molecular features that control site selection, DNA cleavage and recombination.^3–7^

**Figure 1.**
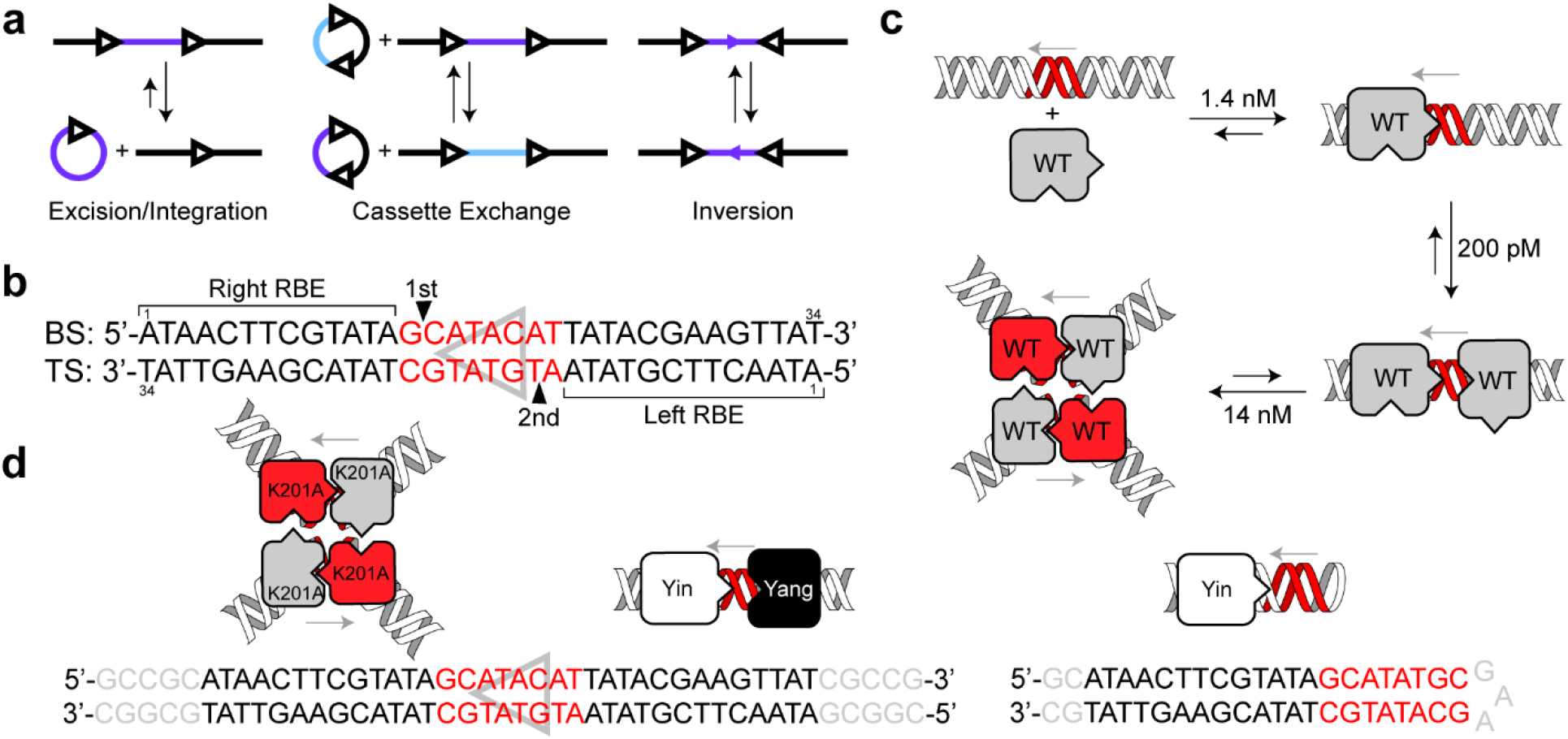
Cre function and assembly intermediates. **(a)** Excision, integration, cassette exchange, and inversion reactions that Cre carries out on DNA containing pairs of *loxP* sites (black triangles) and manipulated DNA regions (blue/purple). The direction of the arrows indicates the orientation of each asymmetric *loxP* site.^1^ **(b)** The *loxP* DNA sequence with conventional nomenclature. The spacer sequence is asymmetric (red, grey triangle) and thus defines the overall orientation of the *loxP* site and determines which are the *left* and *right* RBEs (recombinase binding elements). The *top strand* (TS) runs 5’-3’ from the *left* RBE and has the spacer sequence 5’-ATGTATGC-3’, whereas the *bottom strand* (BS) runs 5’-3’ from the right RBE and has the complementary spacer sequence 5’-GCATACAT-3’. Cre recombines DNA with a bias for cleaving opposing sites on the BS, at the right RBE, with later cleavage of the two TS (filled triangles).^2, 3^ **(c)** Wild type (WT) Cre assembles in a reversible, stepwise, and cooperative manner to form a Cre-*loxP* monomer, then a Cre2-*loxP* dimer, followed by antiparallel dimerization of dimers to form a (Cre2-*loxP*)2 homotetramer with each Cre protomer related by pseudo- four-fold symmetry. WT Cre makes and accepts protein-protein contacts (point, pocket) in a circular fashion in tetrameric complexes. Equilibrium constants for each step were established by analytical ultracentrifugation and mobility shift assays.^4, 5^ **(d)** Tetrameric, dimeric and monomeric Cre complexes used in these studies. Tetrameric complexes were prepared using Cre^K201A^ in complex with a full *loxP* site; additional stabilizing GC ‘clamps’ are shown in grey. Dimeric complexes were assembled using Cre^Yin^ and Cre^Yang^ variants featuring complementary “point” and “pocket” mutations, respectively. The palindromic RBE sequences are black, the asymmetric spacer red, and the stabilizing GC clamps in grey. Monomeric complexes were obtained with Cre^Yin^ bound to a DNA hairpin with a single RBE and a symmetric *loxA* spacer.^6, 7^

### Tetrameric Cre-*loxP* structure

Much of our understanding of the structural bases for site selection, assembly, and recombination of *loxP* DNA by Cre comes from a series of crystal structures of similar tetrameric assemblies;^9, 13–15^ until now, structures of assembly precursors have not been available. Analytical ultra-centrifugation (AUC) and mobility shift assays have shown that Cre is monomeric in solution and cooperatively oligomerizes at nanomolar concentrations in the presence of a *loxP* DNA substrate (Figure 1c).^15, 16^ The available 20+ crystal structures of Cre or Cre-like recombinases in tetrameric complexes exhibit similar characteristics, as exemplified by the tetrameric synaptic pre-cleavage complex between a cleavage-deficient variant, Cre^K201A^, and *loxP* (2HOI^15^;Supplemental Figure 1): (i) The tetramer exhibits pseudo C2-symmetry, in which the *loxP* sites are antiparallel and the protomers bound to the right RBEs are *primed* for DNA cleavage with their tyrosine nucleophile on helix αM positioned within ∼3 Å of the scissile phosphate, while the protomers on the left RBEs are in an *inactive* conformation with their catalytic tyrosines ∼6 Å away from scissile phosphate. In this *BS complex* the *primed* protomers reside on the right RBEs, and the bottom strands (BS) are poised for cleavage. (ii) Each *loxP* DNA duplex is asymmetrically bent by ∼108°. (iii) DNA bending is coincident with extensive protein-protein interactions between protomers on the same duplex, and across the synapse. These interactions include reciprocal cyclic *trans* docking of the C-terminal alpha helix αN and a loop between β strands 2 and 3 (the β2-3 loop) in a manner hypothesized to act as allosteric switches.^13, 17^ This structural asymmetry in the C-terminus and in inter-protomer interfaces observed in all tetrameric synaptic complexes of Cre suggest that protein flexibility is important for regulating protomer activity.^18–24^

### loxP site recognition

The structural basis for *loxP* sequence specificity by Cre has been difficult to decipher. Tetrameric crystal structures have revealed that relatively few of the bases in the RBEs or spacer are directly contacted by amino acid sidechains.^4, 13, 25–28^ Biochemical studies using variant Cre and *loxP* sequences have identified specificity determinants that are not well explained by the contacts observed in tetrameric structures. These observations led to the conclusion that shape complementarity and water-mediated contacts are at least as important as direct readout in sequence selectivity by Cre.^3, 13, 17, 25, 29–31^ Nevertheless, in the absence of high-resolution structures of assembly intermediates, it has remained unclear whether additional protein-DNA contacts might play important roles in initial steps of site recognition that are not retained in the assembled tetrameric intasome. For instance, NMR studies of the catalytic domain of Cre showed that the C-terminal region of the protein containing helix αN docks in a *cis* autoinhibited conformation over the DNA binding site in the absence of DNA, and is displaced upon binding to *loxP* DNA, thereby extending into solution.^32^ The cryo-EM structures of Cre bound to DNAs containing one and two RBEs lend clarity to recognition and progressive assembly of intasome complexes, and emphasize the roles of shape recognition, DNA deformation and protein-protein interactions contributing to specificity.

### Activation of Cre

Control over the DNA cleavage activity of the four assembled Cre protomers is critical to faithful DNA recombination, because activation of adjacent protomers would result in double-strand breaks and failed recombination. The mechanism for this control is particularly intriguing due to the close topological and stereochemical similarity of YSSRs to type IB topoisomerases, whose tyrosine nucleophiles are active to cleave DNA as monomers, and which exhibit low sequence selectivity.^33^ Both classes of enzymes sport highly conserved active sites comprised of conserved arginine and lysine residues that coordinate the scissile base and phosphate in similar manners.^33^ Tetrameric crystal structures of synaptic and Holliday junction intermediates of Cre (and other YSSRs) have invariantly shown pseudo-C2 symmetry with opposing pairs of protomers adopting *inactive* and *primed* conformations, as defined above. Comparison of the *primed* and *inactive* protomers in these structures has drawn attention to the β2-3 loop and C-terminal helices αΜ and αN (Supplemental Figure 1).^13, 25^ Solution NMR studies of the C-terminal domain of Cre showed flexibility in the β2-3, αJ-K and αM-N loops but not for helix αN.^32^ However, the αJ-K loop is involved in direct DNA recognition and does not differ between *primed* and *inactive* protomers in crystals, refocusing attention on the β2-3 loop and αM-αN.

### New insights from Cryo-EM

Cryo-EM structures of the Cre assembly intermediates provide important insights into the mechanism of Cre activation. We designed two mutants, Cre^Yin^ and Cre^Yang^, that when added to a full *loxP* site, in a 1:1:1 fashion, will assemble into a Cre2-*loxP* dimer that maintains a native interface along the *lox* site but prevents interactions at the tetramer interface (Figure 1d). Addition of Cre^Yin^ to a DNA hairpin containing a single RBE allowed us to isolate the Cre-*loxP* monomer complex. Lastly, we assembled a precleavage Cre4-*loxP*2 tetramer using the cleavage deficient Cre^K201A^ variant.^15^ The structures illuminate the critical roles of protein and DNA deformation in enabling site recognition, trans-docking of the αN helix, and for formation of protein-protein contacts involving the key regulatory elements β2-3, αM and αN. These insights simultaneously help us understand site selectivity, controlled activation of alternating protomers, and the basis for the observed bias for strand cleavage order. Additionally, our work highlights the use of conventional defocus methods for high resolution structure determination of low molecular weight macromolecular complexes by cryo-EM.

## Results

### Characterization of isolated Cre assembly intermediates

Wild-type Cre, Cre^K201A^, Cre^Yin^, and Cre^Yang^ were purified to > 95% homogeneity as indicated by band intensities from Coomassie stained SDS-PAGE gels (Figure 2a). WT Cre, Cre^K201A^, Cre^Yin^ (D33A/A36V/R192A), and Cre^Yang^ (R72E/L115D/R199D) were assayed against a linearized plasmid containing two *loxP* sites oriented in parallel (Figure 2b). Cre excises the intervening sequence to produce a circular 1200 bp product and a linear 4000 bp product, which can be resolved on a 1% agarose gel.^34, 35^ Cre recombines the substrate as expected (to ∼65% completion due to the competing reverse reaction), whereas cleavage deficient Cre^K201A^ does not.^15^ In addition, Cre^Yin^, Cre^Yang^, and the Cre^Yin^/Cre^Yang^ combination did not generate recombinant products.

**Figure 2.**
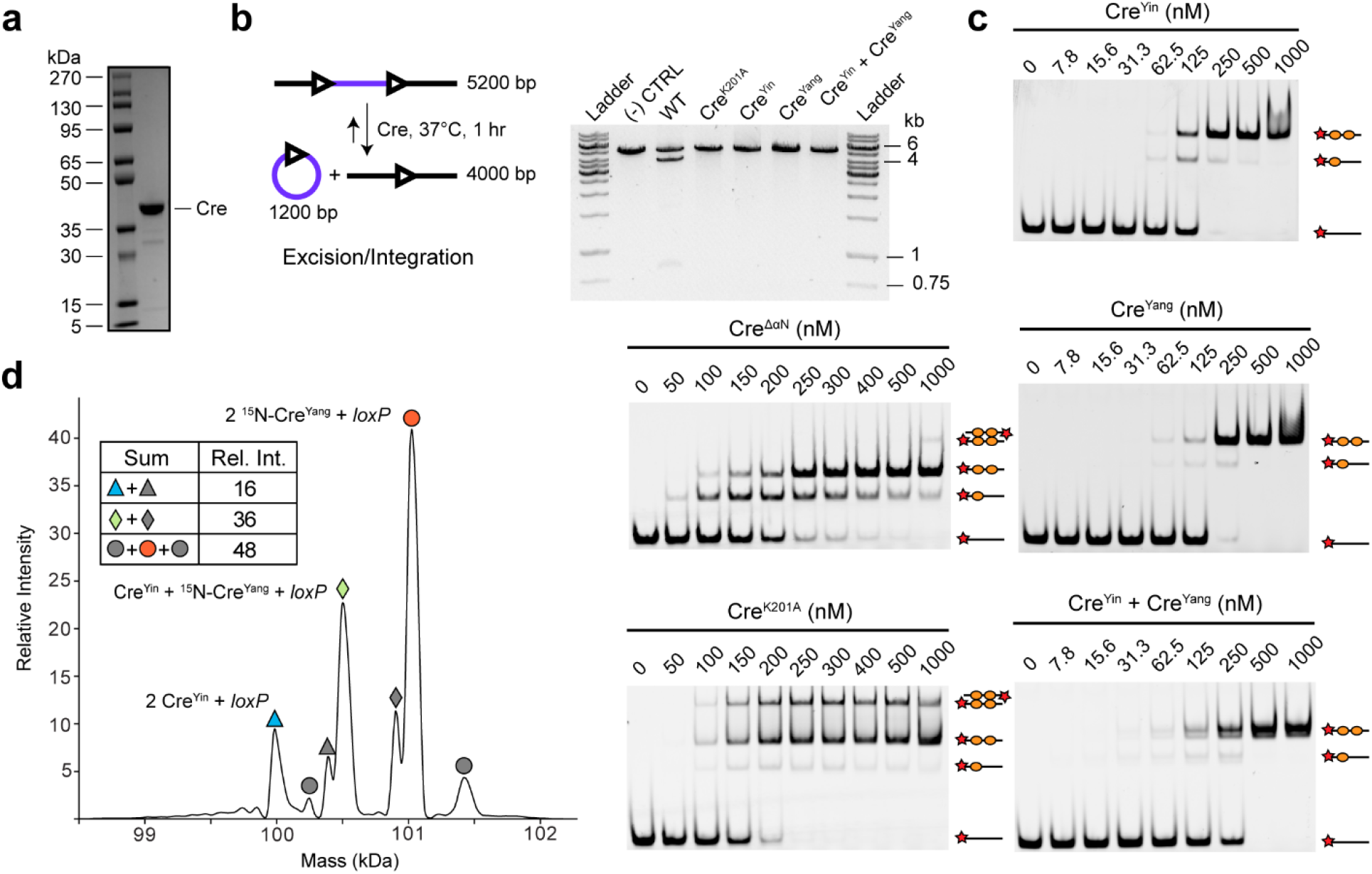
Assembly and recombination of *loxP* DNA by WT Cre and variants Cre^K201A^, Cre^Yin^, Cre^Yang^, and Cre^ΔαN^. **(a)** Representative Coomassie-stained 8-16% SDS-PAGE of SEC purified WT Cre (∼95% purity). **(b)** In vitro recombination assay in which 150 ng of a linearized 5.2 kb plasmid with two parallel *loxP* sites is treated with 500 nM Cre at 37 °C for 1 hr and visualized on a 1% agarose gel. **(c)** Electrophoretic mobility shift assays (EMSA) on 10% polyacrylamide gels, with 10 nM fluorescently labeled *loxP* DNA and varying concentrations of Cre^ΔαN^, Cre^K201A^, Cre^Yin^, Cre^Yang^, or Cre^Yin^ + Cre^Yang^. Unbound DNA and protein- DNA complexes with 1:1, 2:1 and 2:2 stoichiometry are indicated schematically. **(d)** The zero-charge deconvolved ESI mass spectrum of the 2:1 protein:*loxP* complexes formed by Cre^Yin^, [U-^15^N]-Cre^Yang^ and *loxP* DNA. Masses correspond to two Cre^Yin^ + *loxP* (blue triangle), Cre^Yin^ + [U-^15^N]-Cre^Yang^ + *loxP* (green diamond), and two ^15^N- Cre^Yang^ + *loxP* (orange circle). Grey shapes are assigned as proteoforms of these complexes. The summed relative intensities of the three complexes are indicated in the inset table.

We used EMSA to assay the Cre constructs for their ability to assemble with *loxP* DNA at concentrations approximating the equilibrium dissociation constant for synapsis of WT Cre.^15^ In this assay, each Cre mutant was combined with a Cy5-labeled *loxP* DNA and electrophoresed on a 10% native page gel. For Cre^K201A^, which has been shown to be capable of synapsis but not cleavage,^15^ we observed free DNA, monomers, dimers, and tetramers (Figure 2c). Using 10 nM *loxP* DNA and various amounts of Cre^K201A^ we observe about 50% dimers and 50% tetramers, consistent with the published synaptic KD of ∼8 nM; moreover, the monomer to dimer transition is highly cooperative as indicated by a near non-existent population of monomers.^15^ Next, we tested a Cre deletion mutant (ΔαN, residues 1-330) that has been shown to be synapsis-defective.^15^ Upon addition of Cre^ΔαN^ to *loxP* DNA, we can observe that most, if not all, cooperativity is lost, and the formation of tetramers is strongly disfavored (Figure 2c). Cre^Yin^ and Cre^Yang^ formed homodimers in a highly cooperative manner, but no tetrameric species was observed under these conditions (Figure 2c). When Cre^Yin^, Cre^Yang^, and *loxP* DNA were combined, a mixture of species can be observed: the migration distance of the Cre^Yin^/Cre^Yang^ homo- and heterodimers were almost indistinguishable, and this warranted further studies with higher resolution analytical tools.

We first screened each individual component’s mass and purity (Supplemental Figure 3), and subsequently analyzed mixtures of Cre^Yin^, Cre^Yang^ and *loxP* DNA using native mass spectrometry. Native mass spectrometry (nMS) uses soft ionization methods and non-denaturing solution conditions to record gas-phase mass spectra that preserve non-covalent interactions reflective of those in solution.^36^ To increase the mass separation between Cre^Yin^ (38,307 Da) and Cre^Yang^ (38,342 kDa) we used uniformly ^15^N-labeled Cre^Yang^ (38,844 Da, if fully labeled); mass analysis showed > 95% labeling (Supplemental Table 1). The experimentally determined masses of Cre^Yin^ and Cre^Yang^ matched the theoretical masses, but some minor species were present for ^15^N-Cre^Yang^ and Cre^Yin^. These are likely truncations, modifications, or specific salt adducts (Supplemental Figure 3, Supplemental Table 1). The ^15^N-Cre^Yang^ spectrum contains two charge state distributions with the higher charge state centered around +17 being indicative of unfolded or disordered protein.^37^ The determined mass for *loxP-*38 (*loxP* site with a GC clamp on each end) matched the theoretical mass. (Supplemental Figure 3, Supplemental Table 1). In all spectra many of the peaks had additional mass spacing of ∼22 Da attributable to sodium adducts.

Native mass spectra were recorded of samples containing 0.10 µM *loxP* and either 1) 0.50 µM Cre^Yin^, 2) 0.50 µM ^15^N-Cre^Yang^, or 3) 0.25 µM Cre^Yin^ + 0.25 µM ^15^N-Cre^Yang^ in a solution of 200 mM ammonium acetate. For each sample, we observed signals corresponding to the molecular weights of complexes with protein:*loxP* stoichiometries of 1:1 (monomeric complexes) and 2:1 (dimeric complexes)(Supplemental Figure 4). When both Cre^Yin^ and ^15^N-Cre^Yang^ were mixed with *loxP*, we observed abundant signals corresponding to both homo- and heterodimeric complexes with *loxP* DNA (Cre^Yin^-Cre^Yin^, Cre^Yang^-Cre^Yang^, and Cre^Yin^-Cre^Yang^) (Figure 2d). Absent interactions between protomers, we would expect statistical population ratios of 1:1:2 for these complexes, and an additional bias in favor of the heterocomplex due to their complementary protein-protein interfaces. However, the ion counts for the Cre^Yin^ + ^15^N-Cre^Yang^ was only ∼36% of the 2:1 complexes; this may reflect imperfect proportionality between ion counts and solution populations.^38^ Nevertheless, this compositional heterogeneity decreased the number of usable particles in subsequent cryo-EM reconstructions.

### Cryo-EM of Cre assembly intermediates

Three Cre-DNA complexes were examined by Cryo-EM: Cre^Yin^-*loxA* (monomer), Cre^Yin^-Cre^Yang^- *loxP* (dimer), and Cre^K201A^-*loxP* (tetramer) were assembled, purified using size exclusion chromatography, vitrified, and imaged using a Titan Krios G3i microscope (Thermo Fisher Scientific) equipped with a Gatan K3 detector and energy filter. Images were subjected to single particle analysis via 2D classification of motion corrected and CTF estimated particles, followed by 3D classification and non-uniform refinement in cryoSPARC v3.0.^39, 40^ Additionally, 3D Variability Analysis was performed on the tetramer dataset.^41^ This resulted in 3D density maps with global resolutions (gold standard FSC cutoff of 0.143) of 3.91 Å for the Cre^Yin^-loxA monomeric complex (Figure 3a, Supplemental Figure 5, Supplemental Table 3), 4.48 Å for the Cre^Yin^-Cre^Yang^-*loxP* dimeric complex (Figure 3b, Supplemental Figure 6, Supplemental Table 3), and 3.23 Å for the K201A-*loxP* tetrameric complex (Figure 3c, Supplemental Figure 7, Supplemental Table 3). Local resolutions varied from 3.3 Å to 7.1 Å, from 4.1 Å to 7.0 Å, and from 2.9 Å to 5.4 Å, respectively. All single particle reconstructions were performed without the use of symmetry. Atomic models were built in an iterative fashion using Coot, ISOLDE, and Phenix real space refinement.^42–44^ Protein residues 1-19 in all complexes had no resolvable density and were omitted from modeling. Data collection, data processing, and model building statistics are summarized in Supplemental Table 1.

**Figure 3.**
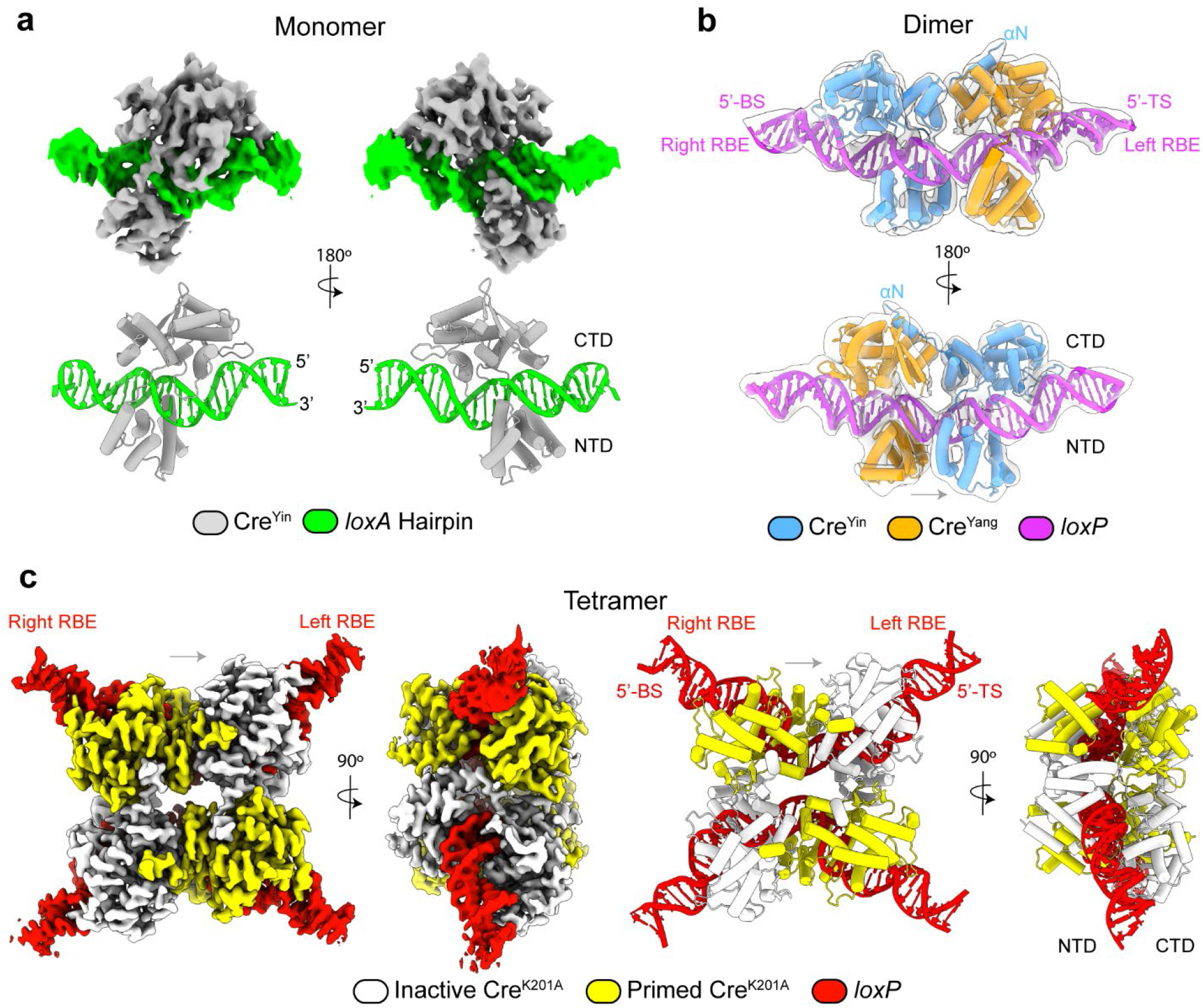
Cryo-EM maps and refined models of Cre-*loxP* assembly intermediates. **(a)** Cre^Yin^-*loxA* half-site DNA complex (threshold of 0.55); hairpin DNA is green and Cre^Yin^ is grey. **(b)** Cre^Yin^-Cre^Yang^-*loxP* hetero- dimer (threshold of 0.5); with Cre^Yin^ in blue, Cre^Yang^ in orange, and the *loxP* DNA in magenta. Cryo-EM map shown in transparent white. The dimer is bent in a direction that favors bottom strand (BS) cleavage, on the right RBE. **(c)** BS tetrameric K201A-*loxP* synaptic complex (threshold of 0.4); with *loxP* DNA in red, protomers in *primed* and *inactive* conformations colored yellow and white, respectively. Grey arrows indicate spacer orientation.

### Cre^Yin^-loxA half-site map and model

The density map for the Cre^Yin^-*loxA* complex contains both domains of the protein, their interdomain linker, and the fully resolved *loxA* hairpin (Figure 3a). Helices αB, αD, and αJ, participants in all the major groove protein-DNA interactions (Supplemental Figure 8), and catalytic core (Supplemental Figure 8) were of the highest resolution (between 3.3 Å and 4.0 Å), allowing for accurate sidechain placement into the map.^45^ Density is present for the β2 and β3 strands that form the anchor for the β2-3 loop, but no robust density was observed for the loop itself (residues 199 - 207, Figure 4a,b), which includes the active site residue K201. Likewise, clear density was observed for αM and the catalytic tyrosine, placing the Oη of Y324 ∼6 Å from the scissile phosphate (Figure 4f), but no density existed to model αN. Thus residues 199-207 and 330-343 were excluded from the model. The *loxA* hairpin phosphate backbone, major, and minor grooves were well resolved, and in the higher resolution regions of the DNA individual ribosyl groups can be visualized. To better visualize the global DNA bend, the *loxA* hairpin model was extended with B-form DNA to contain a second RBE (Figure 5a, monomer, white RBE).^46, 47^ The *loxA* hairpin DNA exhibits an overall bend of 18° with a maximal bend located at the RBE-spacer junction (Figure 5a,c).^48^

**Figure 4.**
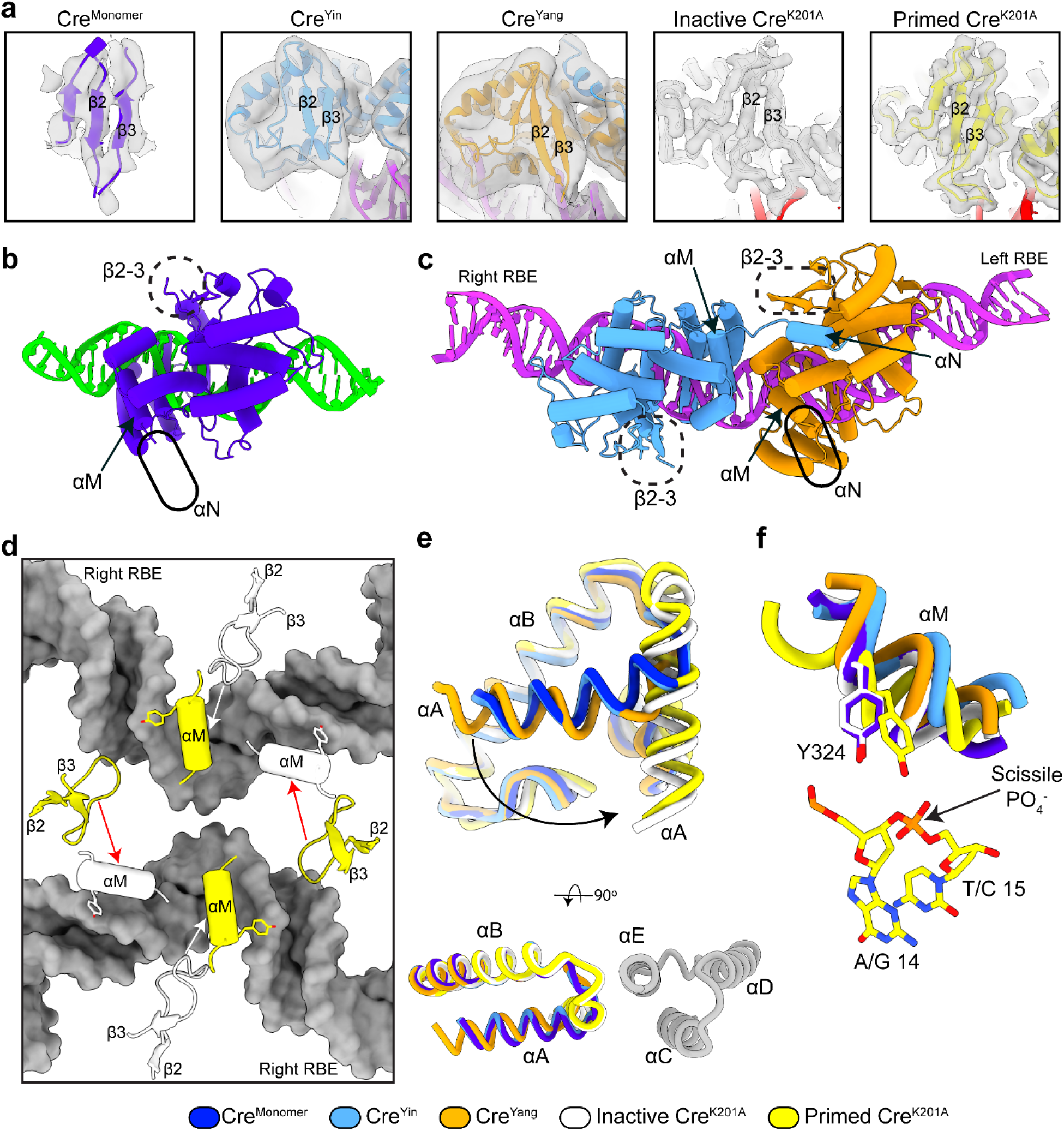
Progressive structural changes that accompany assembly and activation of Cre-*loxP* complexes. **(a)** EM density (grey) and model of the β2-3 hairpin and loop for each protomer; density for the loop is only observed in the tetrameric complex. **(b)** Monomer and **(c)** dimer atomic models with solid and dashed ellipses highlighting missing αN and β2-3 loop density, respectively. **(d)** Intra- and inter- molecular β2-3 contacts help position αM of primed protomers (yellow) in the tetrameric structure. Close contacts (white arrows) result in activation along the duplex; loose contacts (red arrows) are insufficient to activate protomers across the synapse. *loxP* DNA is grey. **(e)** Selected region of superposition from Fig. 5 showing the difference in structure of αA between the monomer, dimer, and tetramer protomer. A second protomer from the tetramer is shown in grey where αA docks against αC and αE of the neighboring protomer. **(f)** Selected region of superposition from Fig. 5 showing αM and Y324 position in contrast to the scissile phosphate for the monomer, dimer, and tetramer protomers.

**Figure 5.**
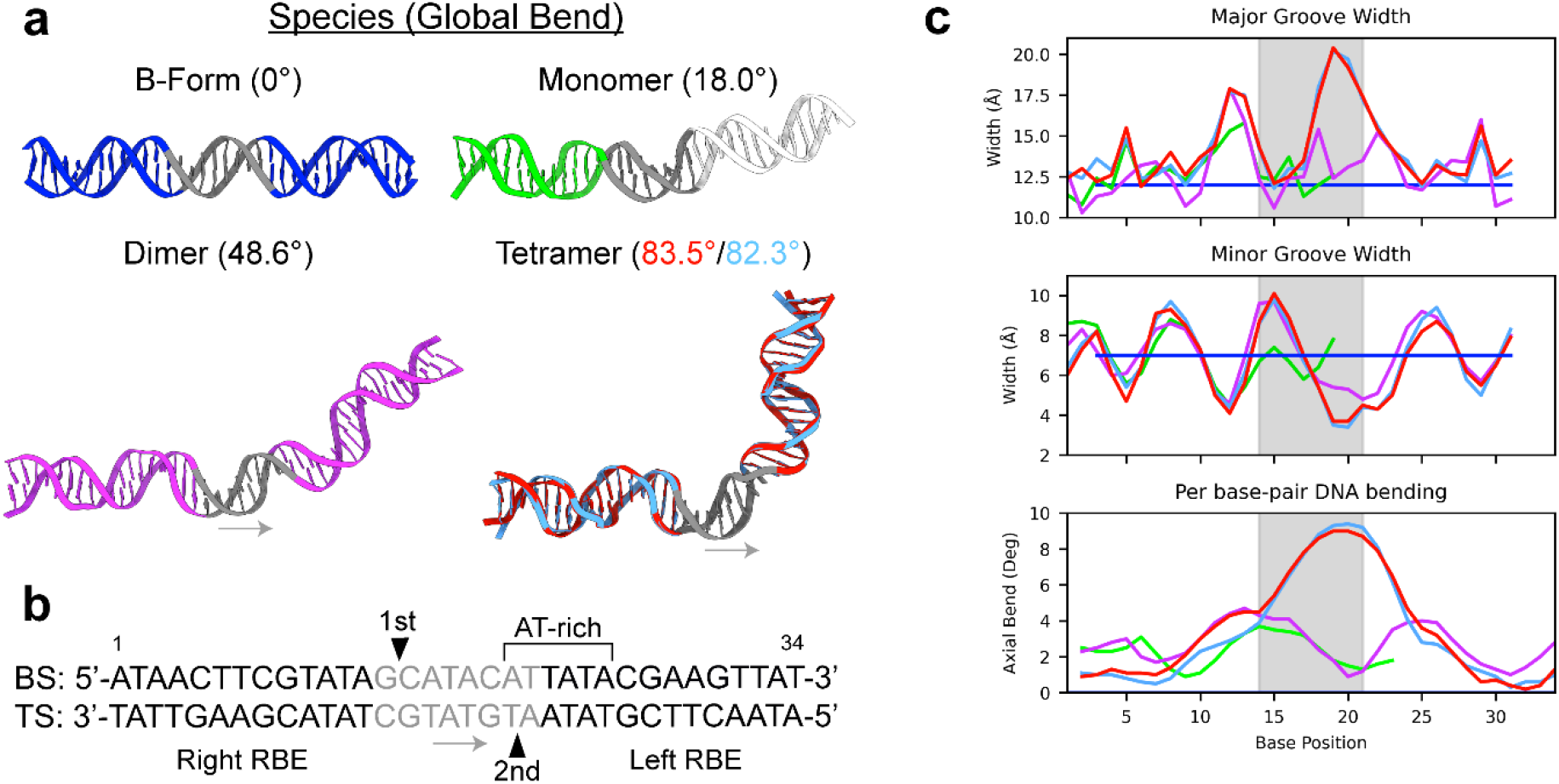
DNA bending and deformations of *loxP* DNA in assembly and recombination intermediates. **(a)** Comparison of global bends for B-form (blue), extended hairpin (green with B-form extended in white), dimer (magenta), and tetrameric (light blue and red; spacers are grey) Cre-*loxP* complexes. Grey arrows indicate spacer orientation. **(b)** *loxP* DNA sequence as shown in **a**. **(c)** Major and minor groove widths and local, per base-pair DNA bending for each *loxP* DNA substrate. Spacer shaded in grey.

### Cre^Yin^-Cre^Yang^-loxP map and model

Single particle analysis of the Cre^Yin^-Cre^Yang^-*loxP* mixture was complicated by the presence of homo- and heterodimers (Cre^Yin^-Cre^Yin^, Cre^Yang^-Cre^Yang^, and Cre^Yin^-Cre^Yang^ on *loxP*), as described above. During 3D classification, three populations of dimers emerged. The resolution of the maps was insufficient to assign the identity of each bound protomer as Cre^Yin^ or Cre^Yang^. In two of the three populations, the N-terminal domain of one protein is seen to be loosely bound and the other is tightly associated with the rest of the complex (Supplemental Figure 6B); these classes were assigned as homodimers on the expectation that engineered electrostatic repulsion would prevent tandem N-terminal domains docking in these species. The relative populations for each dimer in the 3D classification were close to that of native MS (Figure 2d), justifying analysis of the classes with fully bound protomers as the Cre^Yin^-Cre^Yang^-*loxP* BS complex, with Cre^Yin^ bound to the right RBE, donating helix αN to Cre^Yang^ bound to the left RBE. However, at the resolution of the maps, it is also possible that some of the particles featured the opposite arrangement relative to the spacer.

After selecting for fully assembled dimers, it was apparent that this particle set had a third protein loosely bound (Supplemental Figure 6). This third protein was faintly visible in 2D classes and in 3D reconstructions as a smearing of density that originated from the end of helix αM of the density assigned to the Cre^Yang^ protomer; therefore, a particle subtraction routine was used to remove density of the loosely associated protein. Native mass spectra did not reveal a Cre3-*loxP* complex, further supporting a loosely bound third protomer. In the resulting map, we could identify two Cre protomers (Figure 3b, blue, orange) bound to a full *loxP* DNA site (Figure 3b, magenta). Due to the lower resolution of this map, modeling was guided by secondary structural elements in the monomer (above) and tetramer (below). The *loxP* DNA structure was well defined in the map with an asymmetric global bend of ∼48°(Figure 5a,c); this bend is consistent with that measured by phase-sensitive gel mobility shift.^49^ Density is clearly visible for the αN helix of the Cre^Yin^ protomer docked into the αN binding pocket of Cre^Yang^. Compared to crystal structures and the tetramer (below), the αM-αN linker adopts an extended conformation, thereby comprising the only extensive inter-protomer interactions in the dimer complex (Supplemental Table 4). The αN helix of Cre^Yang^ was likely removed via particle subtraction due to its association with a third protein, and therefore, residues 328 – 343 were not modeled. The beta sheet comprised of strands β1, β2, and β3 strands of both protomers could be modeled as there was clear density present, but both Cre^Yin^ and Cre^Yang^ lacked robust density for the β2-3 loops, so residues 199 – 207 were also omitted from modeling (Figure 4a,c).

### Tetrameric Cre^K201A^-*loxP* map and model

Local resolutions ranging from 2.9 Å to 3.9 Å for much of the tetrameric complex allowed for accurate modeling of each protein and the *loxP* DNA (Supplemental Figure 7). Clearly resolved backbone and sidechain density permitted modeling of residues 20 – 341 for all four protomers. Sparse density was present for a few residues preceding residue 20, but lack of clear connectivity prevented us from modeling into it (not shown). The *loxP* DNA constitutes the highest resolution portions of the Cre^K201A^-*loxP* map. For much of the DNA, the density allowed for discrimination of purine vs pyrimidine, and accurate placement of bases was enabled by clear density for the phosphate backbone, ribosyl groups, and the bases themselves. Despite this, it is possible that the tetrameric reconstruction arose from averaging of synapses poised for top strand and bottom strand cleavage (Figure 1); if BS and TS cleavage complexes are overall isomorphous, particle classification with the resolution of acquired data would not be sensitive to nucleotide identity within the spacer sequence.^50, 51^ The tetramer is highly isomorphous to that of the crystallized pre-synaptic complex of Cre^K201A^ bound to *loxP*, with an overall backbone RMSD of ∼2.1 Å, sharing most features of the active presynaptic complex.^15^

As seen in crystal structures of Cre in tetrameric complexes, each protomer bound to the right RBE adopts a *primed* conformation wherein Y324 is positioned close to the scissile phosphate on the BS, and the Y324 of the protomer bound to the left RBE is far from the scissile phosphate on the TS, adopting an *inactive* conformation (Figure 4f). Primed and inactive conformers in the tetramer are related by a pseudo C2 axis, such that each dimer can be seen as exhibiting half-the-sites activity. The density map was refined with no imposed symmetry, and a separate homogenous refinement in cryoSPARC with C2 symmetry did not yield a map with significant improvements in resolution (∼ 0.1 Å). Overlay of each Cre2-*loxP* dimer in the tetramer resulted in an all-atom RMSD of 0.4 Å, similar to that reported in other crystal structures of Cre.^13, 15, 52^ Synapsis results in extensive bending of the DNA to about 83 degrees (Figure 5a, tetramer), most prominent at the right side of the spacer, which features six consecutive AT or TA base pairs (Figure 5c).

### Progressive DNA deformations occur during assembly

Comparative analysis of the assembly intermediates reveals the roles of charge and shape complementarity in recognition of the *loxP* DNA sequence. Protein binding results in marked DNA deformations relative to canonical B-form (Figure 5c).^46, 47^ Each RBE (Figure 5c, non-shaded regions) experiences similar deformations due to protein binding. Superposition of protomer models by α-carbons, excluding the regions of varied structure (αA, αM, αN, and β2-3 loop), yields an overall RMSD of 1.1 Å (Figure 6). In all three structures helices αJ, αB, and αD are inserted into the major groove of the *loxP* DNA site at bases 6-9, 10-12, and 12-15, respectively (Figure 6). In addition, the β4-5 loop is inserted into the minor groove at bases 1-3 and the αJ-K loop is inserted into the minor groove at bases 9-10 (Figure 6). These interactions result in widening and narrowing of the major and minor grooves that is consistent among the assembly intermediates. The major and minor groove widths within the spacer region (Figure 5c, shaded region) vary between assembly intermediates and follow the global DNA bend that progresses along the assembly path. The protomers in the monomer and dimer complexes, and the inactive protomers in the tetramer, bury roughly 2,100 Å^2^ of surface area at their respective protein-DNA interface. The primed protomers in the tetramer make more extensive DNA contacts burying roughly 2,500 Å^2^ of surface area (Figure 5b) due to engagement of β2-3 loop into the minor groove and increased interactions of αE with the spacer region.

**Figure 6.**
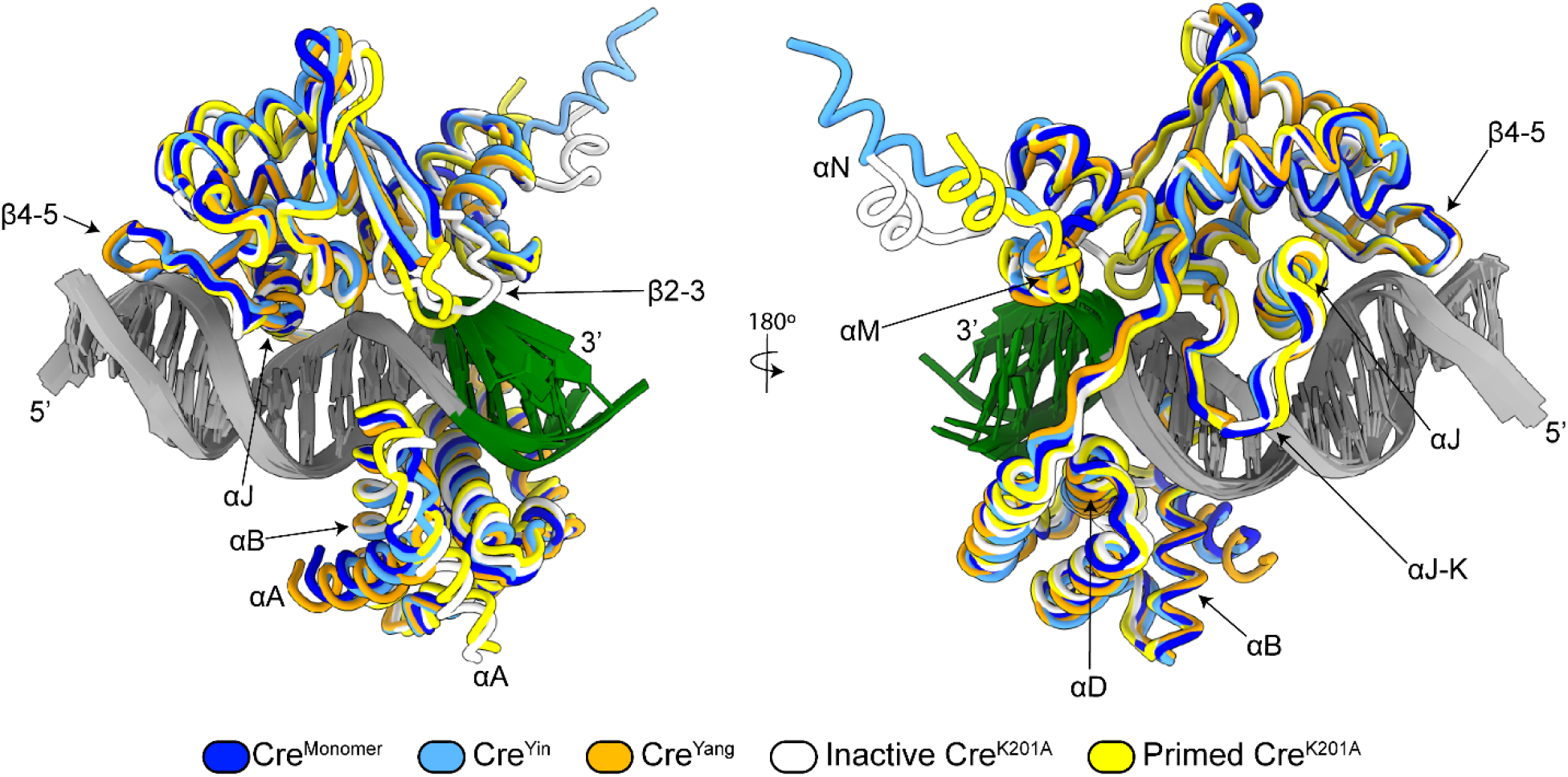
Similarity of site recognition across assembly intermediates. Superposition of the five protein models with important motifs labeled. Superposition performed using all Cαs in α-helices except αA, αM, and αN. *loxP* DNA from monomer structure in translucent grey for reference.

### Evolution of PPIs during assembly of synaptic complexes

Assembly of Cre-*loxP* synaptic complexes is accompanied by formation of protein-protein interactions (PPIs) that position motifs important for DNA cleavage by Cre. In the monomer and dimer structures, there is no visible density for the β2-3 loops of each protomer (Figure 4a,b) and two of the three αN helices (Figure 4b,c). In the dimer, docking of αN from Cre^Yin^ on the right RBE into Cre^Yang^ on the left RBE buries ∼690 Å^2^ of surface area (Supplemental Table 4), but there are no other significant interactions between the protomers. The αN helices and β2-3 loops become structured upon synapsis, with each αΝ helix docking onto neighboring protomers in a cyclic fashion in the tetramer. The β2-3 loops of *primed* protomers on the right RBE insert themselves into the DNA minor groove at the scissile base (Figure 4d). The β2-3 loops originating from the *inactive* protomers on the left RBE are not inserted into the DNA minor groove, and instead pack tightly in a reciprocal fashion against the αM helix of the *primed* protomer whose αN helix is donated to it. Together, the packing of *inactive* protomer β2-3 loops can be seen to confine the αM of each *primed* protomer, while the *primed* protomer does not similarly constrain the αM of *inactive* protomers, allowing those αM-N linkers to adopt extended conformations (Figure 4d, Figure 6). Together with local differences in DNA geometry between the primed and inactive sites, these structural differences position Y324 of αM at the BS scissile phosphate in the *primed* protomer of each right RBE (Figure 4f). Additional structural changes that accompany assembly include the reorientation of αA: in the monomer and dimer structures αA packs against helix αB, whereas it rotates by about 90° to pack against αE of an adjacent protomer when forming the tetrameric complex (Figure 4e).

### Conformational sampling of Cre tetrameric complexes

A range of conformations were revealed in the synaptic Cre^K201A^-*loxP* complexes. 3D variability analysis (3DVA)^41^ was carried out on the tetramer cryo-EM dataset using two principal components, of which PC1 yielded continuous heterogeneity associated with changes in the degree of *loxP* DNA bending (Movie S1). We used ISOLDE to flexibly fit the consensus model into the start and end conformations of PC1 (Figure 7a) to better interpret the structural changes connecting the first and last frames.^44^ The bending angle of the DNA varies by about 10° over the course of the modeled heterogeneity (Figure 7b).^48^ In the first frame (F000), the *loxP* DNA is bent at about 75°, and there is no clear density for either inactive or primed β2-3 loops (Figure 7b,c). The angle of DNA bending increases to about 85° in the last frame (F019) as the protomers rotate along the *loxP* DNA interface, and clear density for the β2-3 loop is observed (Figure 7b,c). Increased DNA bending and rigidification of the β2-3 loop is thus coincident with positioning of the primed protomer’s Y324 near the scissile phosphate (Figure 7d).

**Figure 7.**
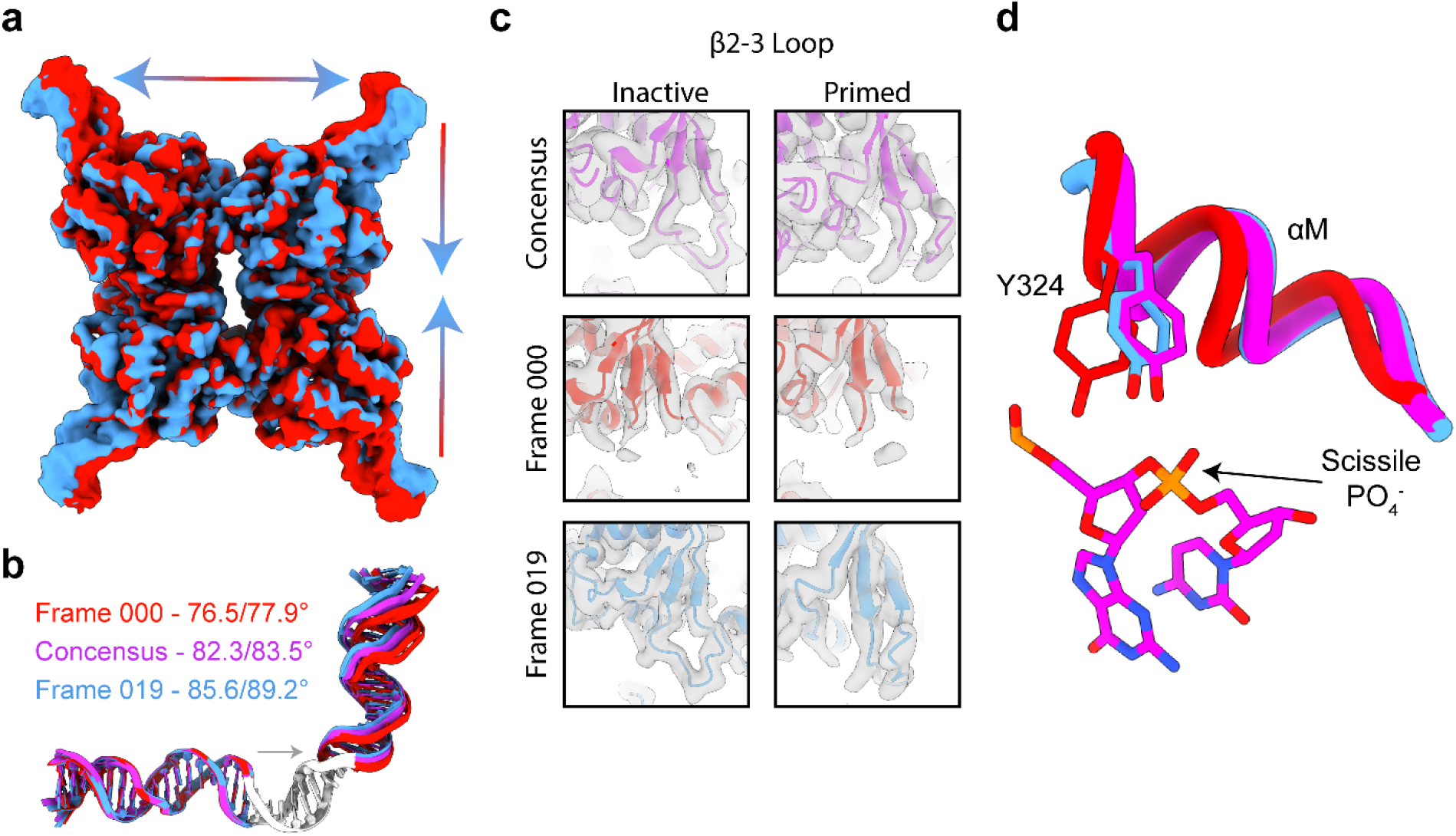
Flexibility in protein-protein interfaces and differing degrees of protomer activation in 3D Variability Analysis of the Cre^K201A^-*loxP* dataset. **(a)** Density maps filtered at 4 Å for frame 000 (red) and frame 019 (sky blue). Arrows indicating the interpolated movement between the start and end frames. **(b)** Superposition of modeled DNAs from frame 000 (red), frame 019 (blue), and the consensus refinement (magenta). **(c)** Density and models for the β2-3 loop for frame 000, frame 019, and consensus refinements. **(d)** Models from consensus (magenta), frame 000 (red) and frame 019 (blue) refinements showing the relationship between Y324 and the scissile phosphate.

## Discussion

### Cryo-EM offers new insights into Cre assembly and function

These cryo-EM studies of Cre-*loxP* assembly intermediates provide new insights into mechanisms of site recognition, half-the-sites activity, strand cleavage order and concerted strand cleavage. The cryo-EM structure of tetrameric Cre^K201A^-*loxP* complex affirms the key attributes of an active pre-synaptic complex revealed by prior crystallographic studies of the Cre^K201A^-*loxP* complex.^15^ The structure of the Cre-*loxP* monomer shows similar protein-DNA contacts to those in tetramer and reveals the structural basis for the absence of topoisomerase activity. The structure of the Cre-*loxP* monomer (54 kDa) shows similar protein-DNA contacts to those of tetrameric proteins and reveals the structural basis for the absence of topoisomerase activity.

### Monomeric Cre specifically binds and recognizes a single RBE but cannot cleave it

To prevent intermolecular protein-protein interactions meditated through helix αN, we assembled complexes of Cre^Yin^ (which bears a mutation at its αN docking site) on a *loxA* hairpin substrate comprising a single RBE and symmetrized “right” spacer (Figure 1). In the cryo-EM density map of the Cre-*loxP* half-site complex, the αJ-K loop is ordered, consistent with a role in DNA binding. However, no density is observed for helix αN and the β2-3 loop, presumably due to the absence of stabilizing protein-protein interactions. This is consistent with their role in mediating protein-protein interactions with neighboring protomers and structural variability in the *inactive* and *primed* conformations. Indeed, the 3.9 Å structure is highly congruent with the *inactive* protomer in the synaptic complex, with an overall backbone RMSD of ∼1 Å (Figure 6), and Y324 is far from the scissile phosphate (Figure 4f). Since K201 resides at the tip of the β2-3 loop, the active site is not fully formed at this step of assembly. The monomeric Cre-*loxP* complex reveals the same major and minor groove protein-DNA contacts observed in protomers of fully assembled synaptic complexes (Supplemental Figure 8), indicating that sequence specific recognition is fully realized upon a single Cre protein binding to an RBE. Analysis of protein-phosphate backbone contacts between the monomer underscores the role of both base-specific contacts and shape recognition in site selection. Thus, the Cryo-EM structure of the monomer is consistent with both the observed site-specific binding and lack of topoisomerase activity (single strand DNA cleavage).^12, 18, 33, 53^

### Half-sites activity and order of strand exchange inferred from pre-synaptic Cre-*loxP* dimers

The use of Cre^Yin^/Cre^Yang^ pairs allowed for isolation and characterization of pre-synaptic Cre2- *loxP* complexes featuring a native protein-protein dimer interface. A defining feature of dimer assembly is *trans* docking by the αN of Cre^Yin^ onto its docking site on Cre^Yang^, accompanied by an increase in DNA bending and increased DNA deformations of bases proximal to and in the spacer (Figure 5a,c). We see that once two Cre protomers are assembled onto a *loxP* site, DNA bending is required to allow the αN of one protomer to dock into its site on the adjacent protomer; only one αN can dock on its neighbor in a dimeric complex. The resolution of the map was insufficient to unambiguously assign the DNA sequence; however, comparison of local DNA bending parameters and major/minor groove deformations to those seen in the monomer and tetramer structures are consistent with a positive axial bend at the AT-rich end of the spacer, producing a dimeric BS complex. (Figure 5c). In a BS complex, the αN of the protomer bound to the BS can reach its docking site on the adjacent protomer positioned on the TS, while the αN of the recipient protomer will remain undocked and extend out into solution in the opposite direction as the DNA bend. Productive synapsis will occur only when two similarly bent dimers encounter each other in an antiparallel orientation^53, 54^, and in the case of synapsis by two BS dimers *only the protomers on the BS can be activated for cleavage* through packing with the β2-3 loops of the adjacent protomers.

In this model, the order of strand cleavage is determined by the probability of a Cre2-*loxP* dimer forming a BS or a TS bend and encountering another similarly bent dimer. Formation of BS-BS synapses would result in bottom strand first cleavage, TS-TS synapses could cleave the top strands, whereas assembly of BS-TS complexes would result in unproductive, or mutagenic recombination events.^55^ Assuming similar efficiency for recombination by either BS-BS or TS-TS synaptic complexes, an ∼80% preference for BS strand first recombination would be explained by a small 15% bias (∼0.6 *kT*) towards formation of BS-bent dimers [0.65 x 0.65 ÷ (0.65 x 0.65 + 0.35 x 0.35) = 78%].^54^ Thus, the structure of the Cre2-*loxP* dimer provides new insights into the structural basis for cooperative DNA binding (by *trans* αN docking), for the observed BS/TS cleavage preferences (spacer bending), and lack of topoisomerase activity due to poorly formed protein-protein interfaces that fail to assemble a complete active site.

### A model for assembly and activation of Cre Recombinase

The new structures allow refinement of a model in which protein and DNA dynamics play important roles in Cre-*loxP* intasome assembly and activation (Figure 8). (1) Sequestration of helix αN over the C-terminal domain DNA binding surface prevents premature oligomerization of Cre protomers, while flexibility in the linker facilitates C-clamp formation by the independently folding N- and C-terminal domains.^32^ (2) Cre binds to an RBE and extends a flexible helix αN into solution. Incomplete formation of the Cre active site due to insufficient protein-protein interactions to stabilize the β2-3 loop and helices αM and αN prevents premature DNA cleavage, while enabling DNA sequence scanning and intersegmental transfer.^56, 57^ The extended αN helix captures a second Cre protomer using a “fly-casting” mechanism. (3) Flexible bending of doubly-bound *loxP* DNA allows alternative sampling of TS and BS synaptic complexes. Pairing of antiparallel duplexes with matching bends can result in synapsis and subsequent recombination, with either BS- or TS-first cleavage. This conformational sampling is enabled by inherent DNA flexibility, as well as ill formed protein-protein interactions in the dimer. In addition, helix αA of the monomeric and dimeric complexes adopts a conformation that is incompatible with PPIs observed in the tetrameric complex. (4) Productive synapsis of two like dimers is accompanied by additional DNA bending and stabilization of protein-protein interfaces by rearrangements of αA, β2-3, and αM-N. The observed structural variability of the tetramer indicates that synapsis is not coincident with activation, suggesting that conformational sampling described by 3DVA likely hints at the steps following DNA cleavage, where stochastic and concerted conformational sampling involving the flexible protein-protein interfaces facilitates strand exchange and the subsequent isomerization step that exchanges *primed* and *inactive* protomers.

**Figure 8.**
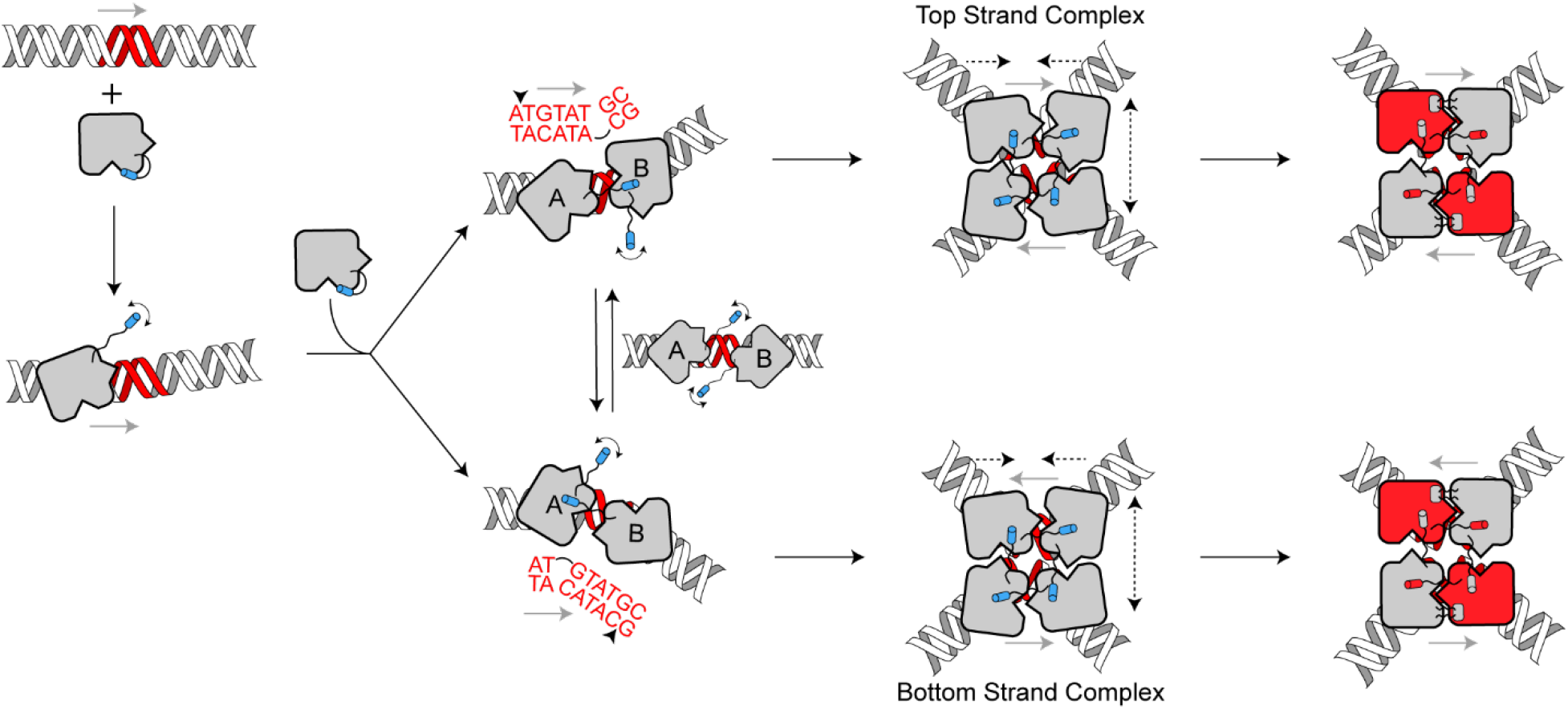
Assembly pathway of Cre recombinase on *loxP* DNA. Binding of the first Cre protomer to an RBE dislodges αN from the autoinhibited state and induces a slight bend in the DNA. A second Cre protein binds to an adjacent RBE, and further DNA bending facilitates intermolecular docking of the αN from one protein into the acceptor pocket of the second protein. Isomerization between top strand and bottom strand complexes can occur through an undocking of αN of one protomer and docking of the other protomers αN. Synapsis of two similarly bent Cre2-*loxP* dimers results in a tetramer that undergoes constrained conformational sampling (arrows) to reach an activated tetrameric complex.

### Implications for developing Cre variants

This work provides new insights into the mechanism of site recognition by Cre. The structure of the Cre-*loxP* RBE monomer features all direct base contacts between DNA recognition motifs (helices αB, αD, αJ, the αJ-K loop, and the β4-5 loop) and the major and minor grooves (Supplemental Figure 8), and the DNA deformations that accompany Cre binding (Figure 5). Additional DNA contacts that coincide with greater DNA bending during the assembly process are limited to phosphate backbone contacts near and within the spacer (Supplemental Figure 8). Since these backbone contacts are not-sequence specific, it is evident that the ability of (Cre2-*loxP*)2 complexes to bend, form the right protein-protein contacts, and form productive active sites are key determinants for productive recombination. 3DVA provides further evidence of this selectivity mechanism wherein DNA bending is coupled to formation of specific productive protein-protein interfaces leading to the positioning of the catalytic tyrosine and activation for DNA cleavage (Figure 7).

Developing Cre variants with altered selectivity requires consideration of multiple factors. Substrate-linked directed evolution experiments against noncognate *loxP*-like DNA sequences that vary in both the RBE and the spacer regions have evolved variants whose mutations fall into two clusters, concentrated at the Cre-*lox* interface or the protomer-protomer interfaces.^4, 6, 58, 59^ Mutations at the Cre-*lox* interface can be understood to alter sequence recognition via formation of complementary protein-DNA interfaces. However, the cluster of mutations in the protein-protein interface cannot be understood in the same context. These Cre variants feature mutations to residues that maps to the protein-protein interface in tetrameric complexes (e.g., D29, M30, R32, Q35, R101, E129, A131, P307, and a duplication of the loop containing G303, V304, and S305). Mutations in the protein-protein interfaces, together with the conformational sampling evident in 3DVA analysis, may relax specificity for formation of cleavage-competent conformations in such Cre-variant tetrameric complexes. In directed evolution experiments, such mutations in early rounds would allow further evolution to maximize efficiency towards the altered *lox* sites. In conclusion, these structures have allowed us to interpret the results of evolved variants of Cre in a structural context, and these results are just the tip of the iceberg for development of alternative Cre-*lox* systems. It is clear to us that there is a need for further experimentation surrounding the influence of protein mutations on the tetrameric complex plasticity and how the per base-pair sequence affects bendability in the spacer region.

## Materials and Methods

### Computational modeling of Cre mutants

In tetrameric pre-synaptic assemblies with pseudo-C4 symmetry, each Cre protomer can be described as having a “point” and a “pocket” that affords specific protein-protein interactions at the interface along the DNA strand and across the synapse (Figure 1c).^13, 15, 52^ Using the crystallographic coordinates of the inactive Cre^K201A^ bound to *loxP* (2HOI) we engineered Cre^Yin^ by mutating the “pocket” of the protein bound to the left side of the *loxP* site and deleted the “point” of the protein bound to the right side of the loxP site to engineer Cre^Yang^ (Figure 1d).^15^ The Proteins, Interfaces, Structures and Assemblies (PISA) tool from PDBe identified residues buried and involved in hydrogen bonding and salt bridge interactions across the synapse at both of the possible interfaces (Fig. S1C, chains A:H and B:G).^60, 61^ Residues D33 and R192 on chains B and H (white protomers) and residues R72 and R119 on chains A and G (black protomers) had the most distinct and expansive hydrogen bonding and salt bridge networks. D33 forms salt bridges with R72 and R119, while R192 forms a salt bridge with E308. We then used the mCSM-PPI2 webserver to perform saturating mutagenesis on the A:H interface along with the B:G interface to understand which mutations of the four residues listed above would provide the most destabilizing effect.^62^ Based on the output of the computational, saturated mutagenesis, we chose to go with the mutations of D33A to eliminate the charge-charge attraction with nearby residues and R192A to disfavor the coordination of αN in the binding pocket for Cre^Yin^. For Cre^Yang^, we flipped the charges on R72E and R119D to create a mostly negatively charged interface and L115D unfavorable charged-hydrophobic interactions with neighboring residues. Although mutation of A36 was not predicted to be detrimental we included the mutation of A36V for Cre^Yin^, in which previously published data shows that this mutation is defective in synapsis (presumably a steric interference, but structurally unverified).^15^ The residues chosen for mutagenesis are shown in in (Supplemental Figure 1).

### Site-Directed Mutagenesis, Protein Expression, and Purification

Mutants were generated using the Q5 Site-Directed Mutagenesis Kit (NEB) in conjunction with a pET-21A plasmid (Novagen) containing WT Cre (provided by Gregory Van Duyne, University of Pennsylvania Philadelphia, PA). Primers were designed manually, are listed in Table S1, and were purchased from Integrated DNA Technologies (Coralville, IA). Each mutant was transformed into E. coli T7 Express LysY (NEB) using electroporation and a 1 mm electroporation cuvette. Transformations were plated on Luria-Bertani (LB) (RPI) media plates supplemented with 100 μg/mL carbenicillin (RPI) and 25 μg/mL chloramphenicol (Fisher Scientific) and incubated overnight (16 hrs.) at 37°C. A single, fresh colony was used to inoculate a 50 mL starter culture of LB supplemented with 100 μg/mL carbenicillin and grown overnight for 16 hours at 37°C in a shaking incubator at 220 rpm. 10 mL of starter culture was used to inoculate a 1L culture of LB supplemented with 100 μg/mL of carbenicillin and left to grow in a shaking incubator at 220 rpm until an O.D.600 of 0.8 was reached (roughly 3 hrs.). Cultures were then induced with 400 μL of 1M IPTG (RPI) and Cre mutants were expressed for 2 hrs. at 37°C and 220 rpm. After incubation, cells were pelleted at 4k rpm and 4°C and frozen at -80°C until purified. [U-^15^N]-Yang was expressed as stated above, but M9 minimal media was used with 1g of ^15^N-ammonium chloride (Cambridge Isotope Labs) was provided as the sole nitrogen source. The expression time was also lengthened to 4 hrs. instead of 2 hrs. due to the slower growing nature of minimal media cultures.

For all proteins (WT, K201A, Yin and Yang) the following purification protocol was followed. Frozen pellets were resuspended on ice in lysis buffer, 40 mM Tris, 300 mM NaCl, 5 mM DTT, pH 7.0 (Fisher Scientific/Fisher Scientific/RPI, respectively). A single complete protease inhibitor tablet (Roche) was added to the cell suspension. The resuspended cells were then lysed using sonication and centrifuged at 15k RPM for 45 min at 4°C. The lysis supernatant was then filtered through a 0.8 – 0.2 um syringe filter (Pall) and diluted with Buffer A (40 mM Tris, 100 mM NaCl, pH 7.0), 1:1. The sample was then loaded onto a SPFF column for cationic exchange purification (GE/Cytiva) and was eluted using a gradient of 100 % buffer A to 100% buffer B (40 mM Tris, 1M NaCl, pH 7.0). Fractions containing the Cre protein were pooled and diluted with buffer A, this time 3:1, buffer A to sample. The diluted sample was then loaded onto a Heparin HP column (GE/Cytiva) for affinity purification and eluted using a gradient of 100% buffer A to 100% buffer B. Fractions containing the Cre protein were combined and concentrated to 1.5 mL using an Amicon Ultra-10, 10kDa MWCO spin filter. The concentrated sample was then loaded onto a GE Superdex 75 prep column and eluted using Cryo-EM buffer 20 mM HEPES, 100 mM NaCl, 5 mM MgCl2, pH 7.0 (RPI/Fisher Scientific/Alfa Aesar, respectively). Fractions containing protein were then pooled and used for further experiments.

### Purification of DNA constructs

ssDNAs were purchased from IDT and shipped dry (Table S1). Upon receipt, ssDNA was dissolved in water to a concentration of 100 μM. Complimentary pairs of dissolved ssDNAs were combined in equimolar amounts. Concentrated Tris base (pH 7.0) and concentrated NaCl were added to each mixture to obtain a final concentration of 10 mM Tris and 100 mM NaCl. Mixtures were then placed into a 95 °C water bath that was shut off and left to cool overnight. After cooling slowly to room temperature, DNA duplex mixtures were loaded onto a QHP ion exchange column (GE/Cytiva) with Buffer A (10 mM Tris (pH 7.0) and 25 mM NaCl) and the duplexes were eluted off with a gradient 0-100% Buffer B (10 mM Tris (pH 7.0) and 1 M NaCl). DNA duplexes eluted near 600 mM NaCl. To concentrate the duplexes and remove salt, samples were EtOH precipitated following a Cold Spring Harbor protocol.^63^

### Purification of Complexes (SEC)

Each protein and DNA complex were formed by addition of 1.2:1 protein to Cre binding sites and were concentrated using an Amicon Ultra-0.5, 3 kDa MWCO spin concentrator yielding 250 μL samples of Cre^Yin^-loxA hairpin, Cre^Yin^-Cre^Yang^-loxP, and Cre^K201A^-loxP at 50 mg/mL, 3.5 mg/mL, and 10 mg/mL. Yin-loxA was purified over a GE Superdex 75 analytical column, and Cre^Yin^-Cre^Yang^-loxP and Cre^K201A^-loxP were purified over a GE Superdex 200 analytical column, both equilibrated with cryo-EM buffer (20 mM HEPES, 100 mM NaCl, 5 mM MgCl2, pH 7.0).

### Recombination Reaction

A 20 μL reaction containing 150 ng of a plasmid containing two loxP sites (Addgene Plasmid #26852) and 500 nM of Cre or Cre mutants in recombination buffer (50 mM Tris-Cl, 50 mM NaCl, 10 mM MgCl2, 1 mM DTT, pH 7.0) at 37 °C for 1 hr.^15, 64^ Reactions are then subjected to a 70 °C water bath for 5 minutes to deactivate Cre. The samples were then combined with 6x gel loading dye (NEB) and ran on a 1%, 10 cm TBE/agarose gel (Fisher Scientific) that was impregnated with SYBR safe (Invitrogen) at 120 V for 55 minutes.

### Electrophoretic mobility shift assay

Gel polymerization and gel running buffers were of the same composition (50 mM HEPES and 20 mM Tris, pH 7.0). 6% native PAGE gels were prepared by first making a gel solution containing 6% acrylamide (29:1) (Fisher Scientific).^15^ This solution was then degassed under vacuum and mixing. 12 mL of degassed solution was mixed with 200 μL of 10% (w/v) ammonium persulfate (Biorad) dissolved in water and 10 μL of TEMED (Acros) and then two 10- well mini gels were poured.

A set of 10 μL reactions containing varying amounts of Cre or Cre mutants (1 – 1000 nM) and 10 nM Cy5-labeled loxP DNA (IDT), were prepared in binding buffer (20 mM HEPES, 150 mM NaCl, 5 mM MgCl2, 2 mM DTT, pH 7.0) along with 50 μg/mL salmon sperm DNA (Invitrogen) and 100 μg/mL BSA (Thermo Scientific).^15, 64^ Binding reactions were equilibrated at room temperature for 30 minutes. 1 μL of 10x EMSA gel loading dye (60% glycerol, 40% binding buffer, 0.1% bromophenol blue) was added to each reaction and then loaded onto a pre-run (120 V, 30 min) 6% native PAGE gel using gel running buffer. Samples were ran at 80 V for 75 minutes and imaged on an Amersham Typhoon Gel imager.

### Native Mass Spectrometry

Native MS experiments were performed on a Q Exactive Extended Mass Range (EMR) mass spectrometer (Thermo Fisher) that was modified to allow for surface-induced dissociation (SID, not used in this work).^65^ Stock solutions of the proteins and DNA were individually dialyzed into 200 mM ammonium acetate (pH unadjusted) using a Pierce 96-well microdialysis plate, 3.5K MWCO (Themo Fisher). The concentrations of the dialyzed stock samples were determined using a NanoDrop 2000/2000c spectrophotometer (Thermo Fisher) and their calculated extinction coefficients. Extinction coefficients for the proteins were calculated using the ExPASy ProtParam tool and the DNA extinction coefficient was calculated using the tool built into the NanoDrop software.^66^ For preparation of the protein-DNA complexes, the stock samples were mixed to their final experimental concentration. Each sample was then injected into an in-house pulled borosilicate filament capillary (OD 1.0 mm, ID 0.78 mm) and subsequently ionized by nano-electrospray ionization in positive mode. Ion optics (including SID device) were tuned to allow for transfer of complex ions to the mass analyzer without causing dissociation. Varying HCD voltage was tested and then set to 60 V for optimal transmission and de-adducting of the complex ions. Other parameters include the following: Spray voltage adjusted to 0.6 kV and then held constant, capillary temperature 250 °C, trap gas pressure set shifted the UHV sensor up to ∼8E-10 mbar, ion inject time 200 ms, averaged micro scans 5, and resolution 8750 as defined at 200 m/z.

All data were manually examined and deconvolved using UniDec V4.4.^67^ UniDec parameters include charge range of 5 to 25, mass range of 10,000 to 1,100,000 Da, sample mass every 1 Da, peak FWHM 0.85 Th, Gaussian peak shape function, beta 50 (artifact suppression), charge smooth width 1, point smooth width 10, native charge offset range -5 to +5, and peak detection range 40 Da. These setting worked well, but manual mode was used to force charge states for known *m/z* regions. This was beneficial for artifact suppression, particularly for optimizing parameters for a range of species present across the *m/z* range. The resulting deconvolutions were plotted as the sum normalized relative signal intensities in the form of zero-charge mass spectra.

### Cryo-EM Grid Preparation and Data Collection

All samples were SEC purified before grid preparation. Before plunge freezing of grids, octyl-n-glucoside was added to a concentration of 16.8 mM to the Cre^Yin^-Cre^Yang^-LoxP and Cre^K201A^ samples, while CHAPSO was added to a concentration of 4 mM to the Cre^Yin^-loxA hairpin sample. 3 μL of Cre^Yin^-loxA hairpin (7.5 mg/mL), Cre^Yin^- Cre^Yang^-loxP (5 mg/mL), and Cre^K201A^- loxP (4 mg/mL) were applied to glow discharged (20 mA, 1 min, Pelco easiGlow) Au Quantifoil R1.2/1.3 300 mesh grids (Ted Pella) and incubated for 60 seconds horizontally. The grids were then loaded into Vitrobot Mark IV (Thermo Fisher Scientific) operating at 4 °C and 100% humidity and blotted for 4s with a blot force of 1 before being plunge frozen into a mixture of liquid ethane/propane.

All data were acquired on a Titan Krios G3i operated at 300 keV with a Cs corrector in nanoprobe EFTEM mode (50 μm C2 aperture, 100 μm objective aperture). A Gatan K3 camera operating in correlated double sampling mode and a BioContinuum energy filter (15 eV slit width, zero-energy loss) were used to record 45 frame movies with a total dose of 60 e^-^/Å^2^. 3,538 movies with two shots per hole were collected for the Cre^Yin^-loxA hairpin sample using a defocus range of -1.0 μm to -2.2 μm at a nominal magnification of 105,000x (0.702 Å physical pixel size). For the Cre^Yin^ - Cre^Yang^-loxP sample, 1,527 movies were collected with a tilt of 28° using a defocus range of -1.5 μm to -3.0 μm, and 2,779 movies were collected with no tilt using a defocus range of -1.0 μm to -2.2 μm at a nominal magnification of 81,000x (0.899 Å physical pixel size). For the Cre^K201A^-loxP sample, 906 movies were collected with a tilt of 28° using a defocus range of -1.5 μm to -3.0 μm, and 2,756 movies were collected with no tilt using a defocus range of -1.0 μm to -2.2 μm at a nominal magnification of 81,000x (0.899 Å physical pixel size). An additional 3,330 movies were collected of the Cre^Yin^-loxA complex to obtain more particles for resolution improvement. These were collected with the same settings listed above.

### Cryo-EM Data Processing

All data processing was carried out using CryoSPARC v3.0.^39^

#### Cre^Yin^-loxA Hairpin

3,538 movies were patch motion corrected and their CTF parameters were obtained using patch CTF estimation. Micrographs were curated by CTF fits less than 4 Å and total motion less than 200 pixels, resulting in a stack of 3,211 micrographs. 300 micrographs were blob picked and 2D classified to yield a set of 47,291 particles that were used to train a TOPAZ model. This model was then used to pick all 3,211 micrographs, yielding 477,224 particles.^68^ Particles were extracted using a 256 x 256 box subsequently down sampled to 128 x 128 (1.404 Å/pixel). Initial 2D classification removing false picks or poorly defined classes left 415,307 particles. These particles were then subjected to ab initio reconstruction using two classes, of which 284,516 particles were left in the class resembling a monomer bound to a DNA hairpin. These particles were then subjected to three rounds of 3D classification that resulted in a stack of 90,940 particles that upon non-uniform refinement resulted in a map with a global resolution of 4.26 Å. Re-centering and re-extraction with a box size of 350 x 350 pixels and down sampled to 256 x 256 pixels (0.959 Å/px) followed by non-uniform refinement resulted in a map with a 4.10 Å resolution using 87,515 particles.^69^ Additionally, 3,330 movies were collected and processed in the same manner yielding 59,200 more particles. In total 146,715 particles were used for the reconstruction of the Yin-loxA hairpin complex, resulting in a final global resolution of 3.9 Å.

#### Cre^Yin^*-* Cre^Yang^*-LoxP*

1,527 28°-tilt and 2,779 0°-tilt movies were independently patch motion corrected and their CTF parameters were obtained using patch CTF estimation. Micrographs were curated by CTF fits less than 4 Å and total motion less than 200 pixels, resulting in a stack of 1,198 and 2,650 micrographs, respectively. Blob picking of 300 micrographs resulted in 62,749 particles and these particles were used to generate templates for template picking. Template picking all micrographs yielded roughly 700k particles. Concurrently, a TOPAZ model was trained on the same 62k, blob picked particles and the model was used to pick all micrographs resulting in 1.9M particles.^68^ These particle stacks were combined, and duplicates were removed to yield a stack of 2.2M particles. Particles were extracted using a 256 x 256 box subsequently downsampled to 128 x 128 (1.798 Å/pixel). 1.2M particles were then subjected to ab initio reconstruction using three classes. Multiple rounds of 3D classification and non-uniform refinement resulted in a stack of 256,734 particles. Particle subtraction of a loosely bound third protein followed by local, non-uniform refinement yielded a map at 4.48 Å.^69^

#### Cre^K201A^*-loxP*

906 28°-tilt and 2,756 0°-tilt movies were independently patch motion corrected and their CTF parameters were obtained using patch CTF estimation. 28°-tilt micrographs were curated by CTF fits less than 4 Å and total motion less than 200 pixels, yielding 806 micrographs. Blob picking of 150 micrographs resulted in 22,250 particles and these particles were used to train a TOPAZ model. The TOPAZ model was used to pick all micrographs, resulting in 1.2M picked particles.^68^ Blob picking of the individual datasets yielded roughly 588k particles. TOPAZ and blob picked particle sets were combined and duplicates were removed to yield 737,980 particles. 2D classification to remove poorly aligning particles resulted in 608k particles that were further processed using 3D methods. Particles were extracted using a 256 x 256 box (0.899 Å/pixel). 608K particles were then subjected to ab initio reconstruction using a single class. A single round of 3D classification with two classes resulted in 315,574 particles that refined to 3.23 Å. These 315K particles were then subjected to 3D Variability Analysis with two modes.^41^

### Model Building

#### Cre^Yin^-loxA Hairpin

Chain A of PDB 2HOI was used as the starting model for the monomer and appropriate mutations were made within PyMol. The hairpin DNA was generated using 3DNA (http://web.x3dna.org/) by compositing a B-form loxA RBE and spacer sequence with a GAA hairpin from PDB 1JVE.^46, 70^ The loxA hairpin and protein model were rigidly docked into the map using ChimeraX.^71, 72^ The complex was subjected to all-atom refinement in Coot using user-defined restraints (all atoms within 5 Å of one another).^42^ Iterative rounds of flexible fitting using the program ISOLDE, implemented in ChimeraX, and real space refinement in Phenix, were used to obtain the final monomer model.^43, 44^

#### Cre^Yin^*-* Cre^Yang^*-LoxP*

The protein chain from the Yin-loxA model was used as the starting model for each protein in the dimer and appropriate mutations were made to each protein chain within PyMol. 3DNA was used to generate a B-form loxP site that matched our DNA construct.^46^ Each DNA and protein model were rigidly docked into the map using ChimeraX.^71, 72^ The complex was then all-atom refined in Coot using user-defined restraints (all atoms within 5Å of one another).^42^ Iterative rounds of flexible fitting using the program ISOLDE, implemented in ChimeraX, and real space refinement in Phenix, were used to obtain the final dimer model. ^43, 44, 71, 72^

#### Cre^K201A^*-loxP*

PDB 2HOI was used as the starting model and was rigidly docked into the map using ChimeraX.^15, 71, 72^ The DNA ends of each strand were extended using PyMol to match the DNA used in the study. Iterative rounds of flexible fitting using the program ISOLDE, implemented in ChimeraX, and real space refinement in Phenix were used to obtain the final tetramer model.^43, 44^

3DVA was performed in CryoSPARC using two modes, a filter resolution of 4.0 A was used and displayed in 20 frames.^41^ ISOLDE was used to fit the consensus model into the first and the last frame output from the 3DVA job.^44^

## Acknowledgements

This work was supported by NIH grants R01 GM122432-04 (MPF), P41 GM128577 (VHW), and S10OD023582. We thank Dr. Greg Van Duyne for providing Cre expression vectors and encouragement and Dr. Aparna Unnikrishnan for preparation of Cre samples. Electron microscopy was performed at the Center for Electron Microscopy and Analysis (CEMAS) at The Ohio State University. We thank Dr. Yoshie Narui (CEMAS) for training and support with Cryo-EM experiments and access to computational resources. Lastly, we thank members of the Foster and Wysocki labs for useful discussions.

---

**Supplemental Figure 1.**
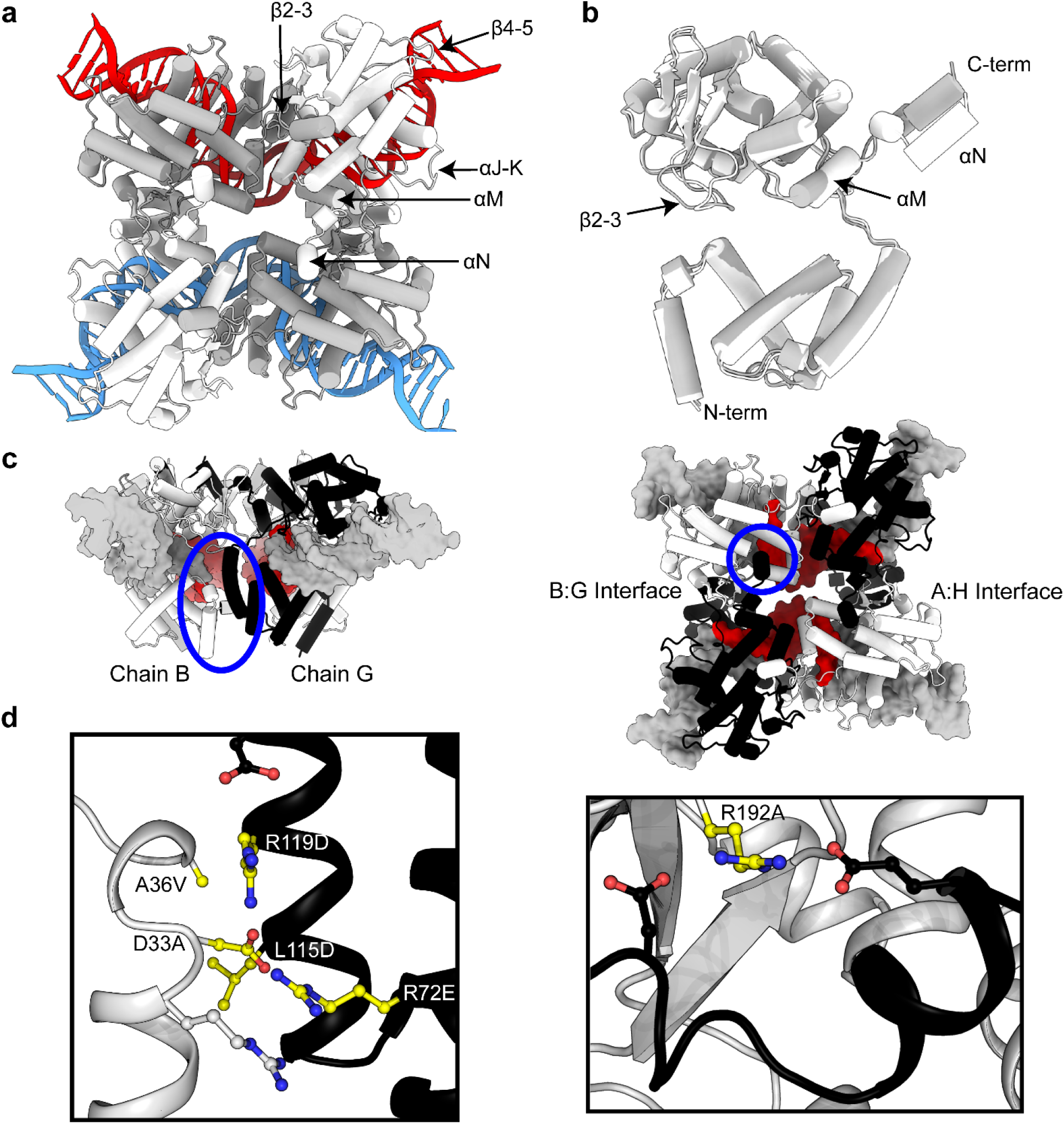
Cre motifs and Yin-Yang mutant design. **(a)** Cylindrical representation of primed protomers (grey) and inactive protomers (white) bound to a surface representation of *loxP* DNA rendered from the Cre^K201A^-*loxP* crystal structure 2HOI.^4^ (**b**) Comparison of primed and inactive protomers showing little structural variability except for αM, αN, and β2-3 loop. Superposition performed using all Cαs in α- helices except αA, αM, and αN. **(c)** Presynaptic Cre^K201A^-loxP crystal structure (2HOI) highlighting the regions chosen to mutate for Yin (white, chain B) and Yang (black, chain G) that include a network of extensive salt bridges and hydrogens bonds between the two N-terminal domains and the αN binding pocket on Yin. **(d)** Close-up of interactions in 2HOI of residues mutated (yellow) to generate Cre^Yin^ (D33A/A36V/R192A) and Cre^Yang^ (R72E/L115D/R199D).

**Supplemental Figure 2.**
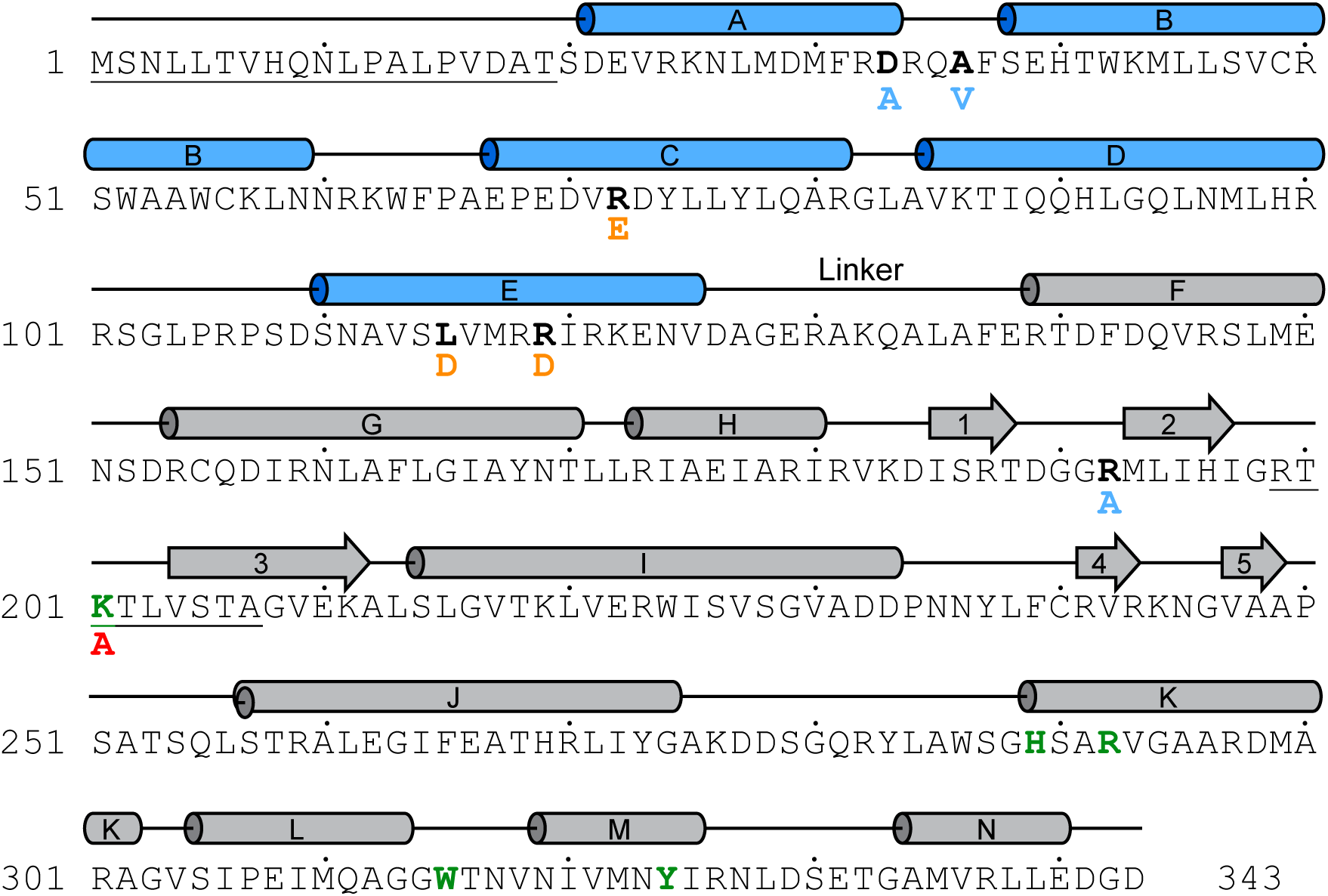
Secondary structure map for Cre recombinase. The N-terminal core binding domain is represented by blue cylinders and the C-terminal catalytic domain as grey cylinders and arrows. Residues in bold black were mutated to obtain Cre^Yin^ (light blue) and Cre^Yang^ (orange). Bold residues in green comprise the active site and Cre^K201A^ mutant shown in bold red. Underlined residues 1- 19 are missing density in all three determined structures and underlined residues 199-207 are missing in the monomer and dimer structure. Dots indicate every 10^th^ residue.

**Supplemental Figure 3.**
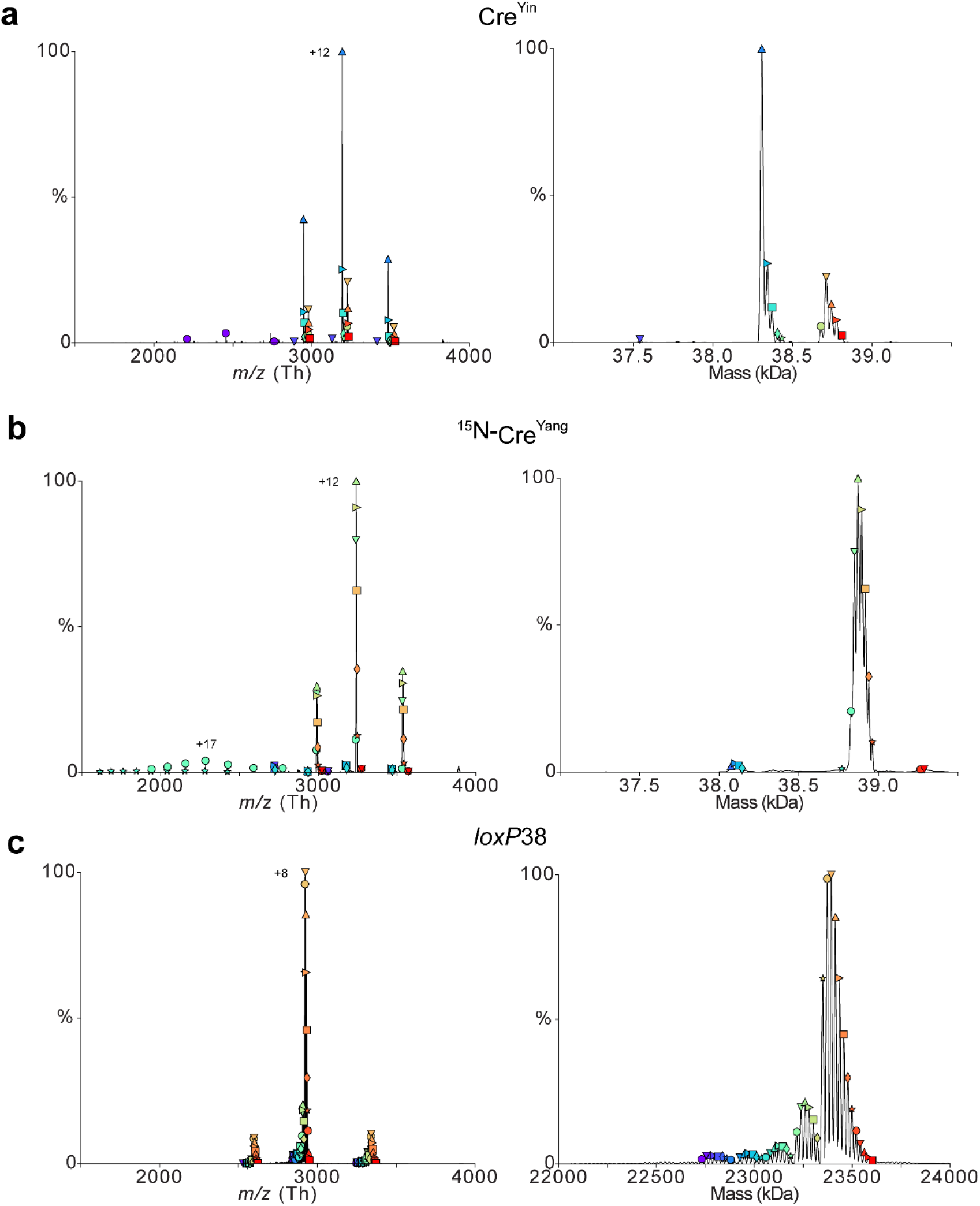
Electrospray mass spectrum (left) and deconvolved zero-charge mass spectrum (right) for **(a)** [U-^15^N]-Yang (5uM), **(b)** Yin (5uM), and **(c)** *loxP*38 (1uM). The *loxP* spectrum features repeating masses in each peak cluster of ∼22 Da assigned to sodium adducts.

**Supplemental Figure 4.**
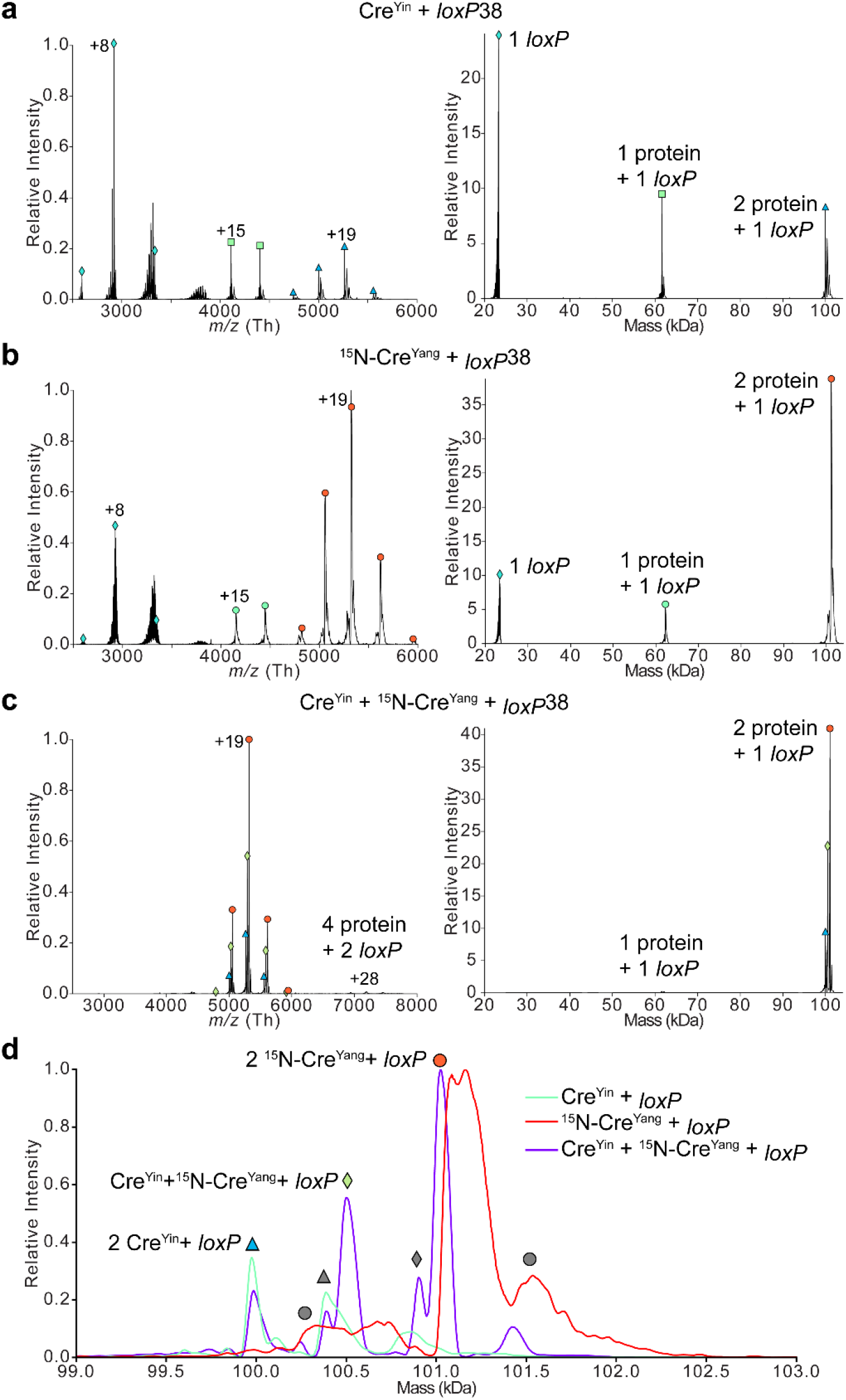
Mass spectrum (left) and deconvolved zero-charge mass spectrum (right) of **(a)** 0.50 µM Yin + 0.1 µM *loxP*38, **(b)** 0.50 µM ^15^N-Yang + 0.1 µM *loxP*38, and **(c)** 0.50 µM Yin + 0.50 µM ^15^N-Yang + 0.1 µM *loxP*38. **(d)** Zero-charge mass spectrum overlay of the three different mixtures zoomed into the 2:1 protein:*loxP* complex mass region. The ^15^N-Yang + *loxP* mixture was not as successfully de-adducted as the other mixtures resulting in the peaks appearing shifted and broader.

**Supplemental Figure 5.**
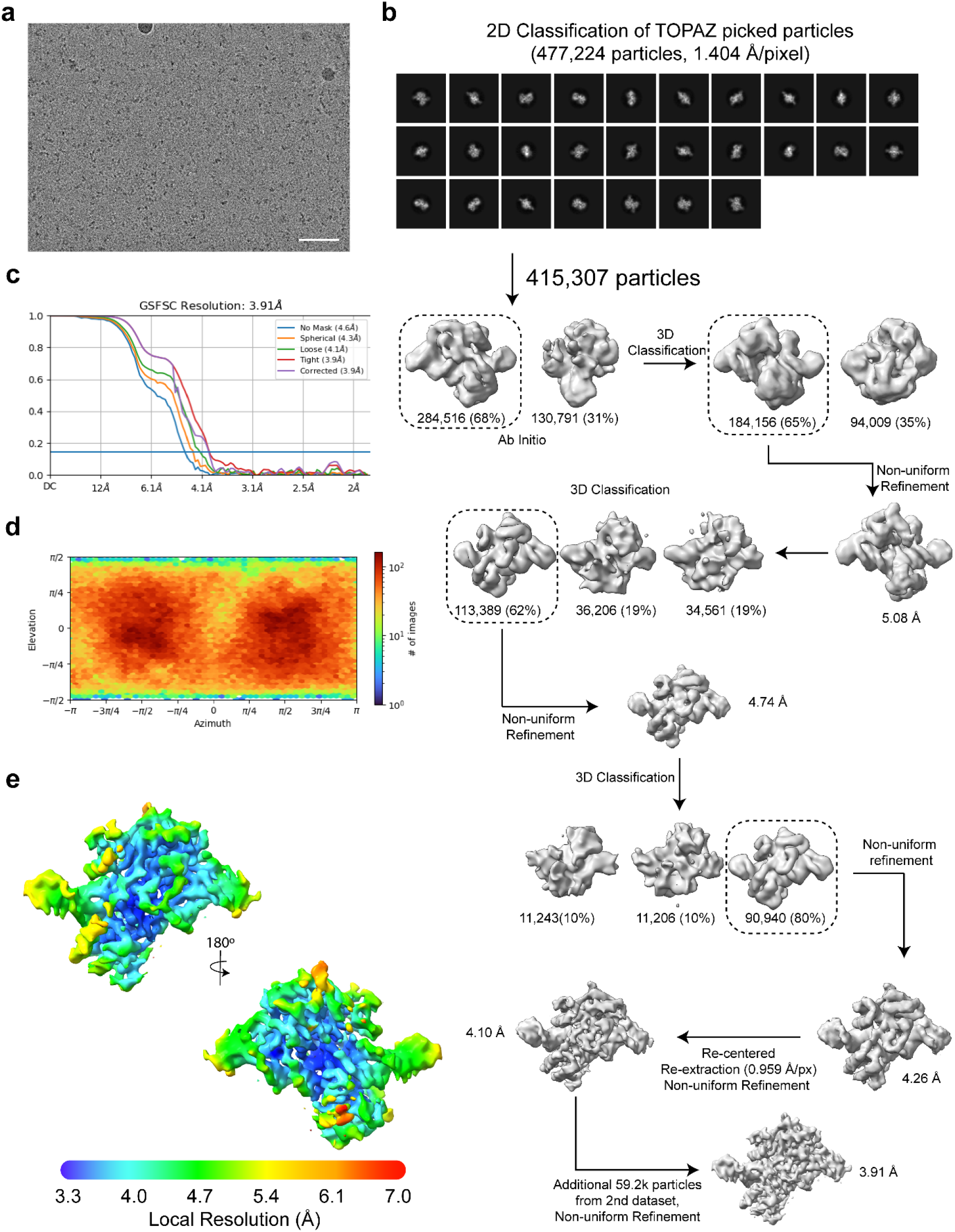
Processing of Cre^Yin^-loxA hairpin. **(a)** Representative micrograph at -2.4 µm defocus. Scale bar indicates 500 Å. **(b)** 2D and 3D processing scheme utilized in CryoSPARC. **(c)** Gold standard FSC plot with global resolution computed at 0.143. **(d)** Orientation distribution for final reconstruction. **(e)** Local resolution plot varying from high resolution (dark blue) to low resolution (red).

**Supplemental Figure 6.**
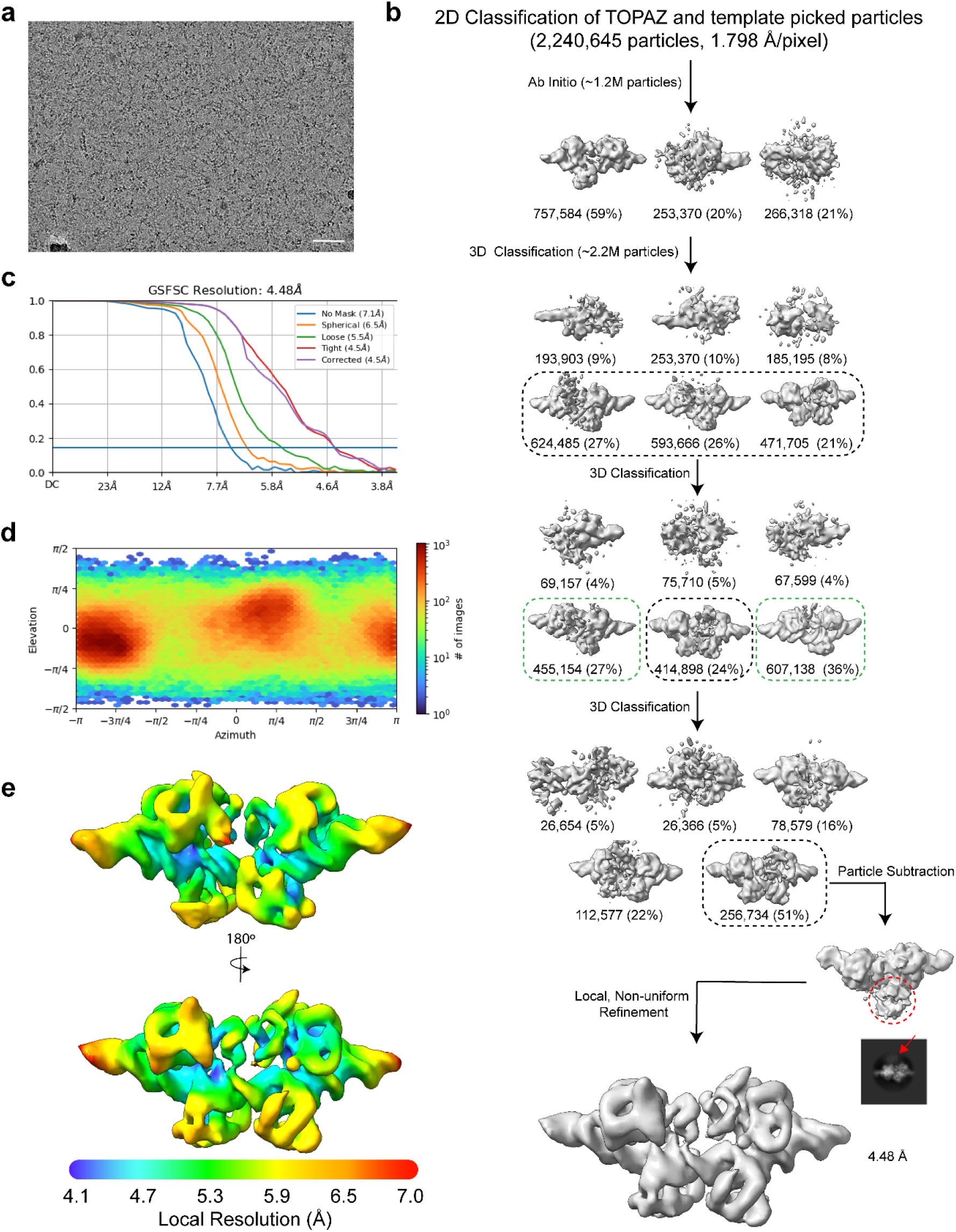
Processing of Cre^Yin^-Cre^Yang^-loxP. **(a)** Representative micrograph at -2.2 µm defocus. Scale bar indicates 500 Å **(b)** 3D processing scheme utilized in CryoSPARC. Combined classes in black rectangle and homodimers with non-interacting core binding domains in green rectangles. Red circle/arrow indicating loosely bound third protein that was selected and removed using particle subtraction routines. **(c)** Gold standard FSC plot with global resolution computed at 0.143. **(d)** Orientation distribution for final reconstruction. **(e)** Local resolution plot varying from high resolution (dark blue) to low resolution (red).

**Supplemental Figure 7.**
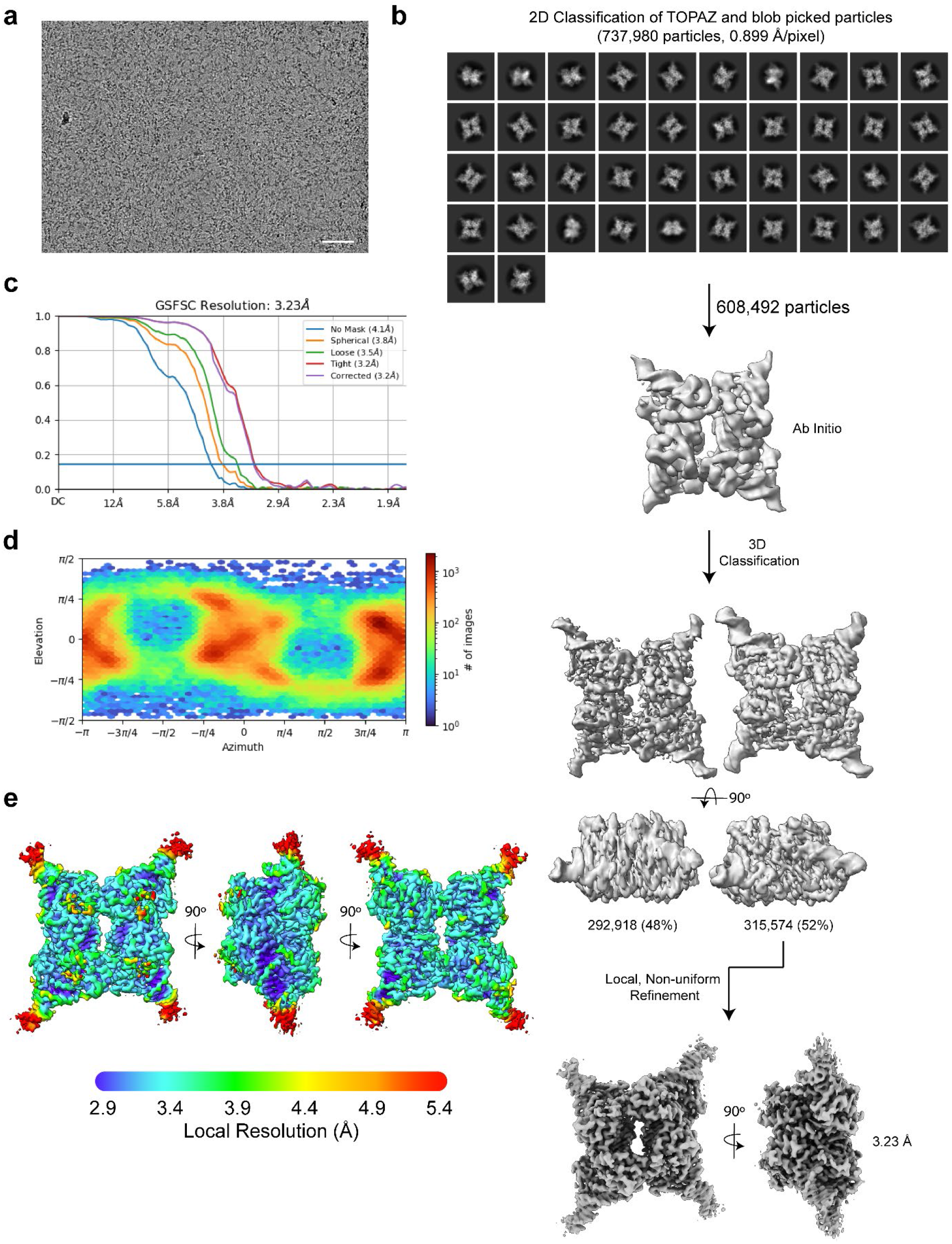
Processing of Cre^K201A^-loxP. **(a)** Representative micrograph at -2.2 um defocus. Scale bar indicates 500 Å. **(b)** 2D and 3D processing scheme utilized in CryoSPARC. **(c)** Gold standard FSC plot with global resolution computed at 0.143. **(d)** Orientation distribution for final reconstruction. **(e)** Local resolution plot varying from high resolution (dark blue) to low resolution (red).

**Supplemental Figure 8.**
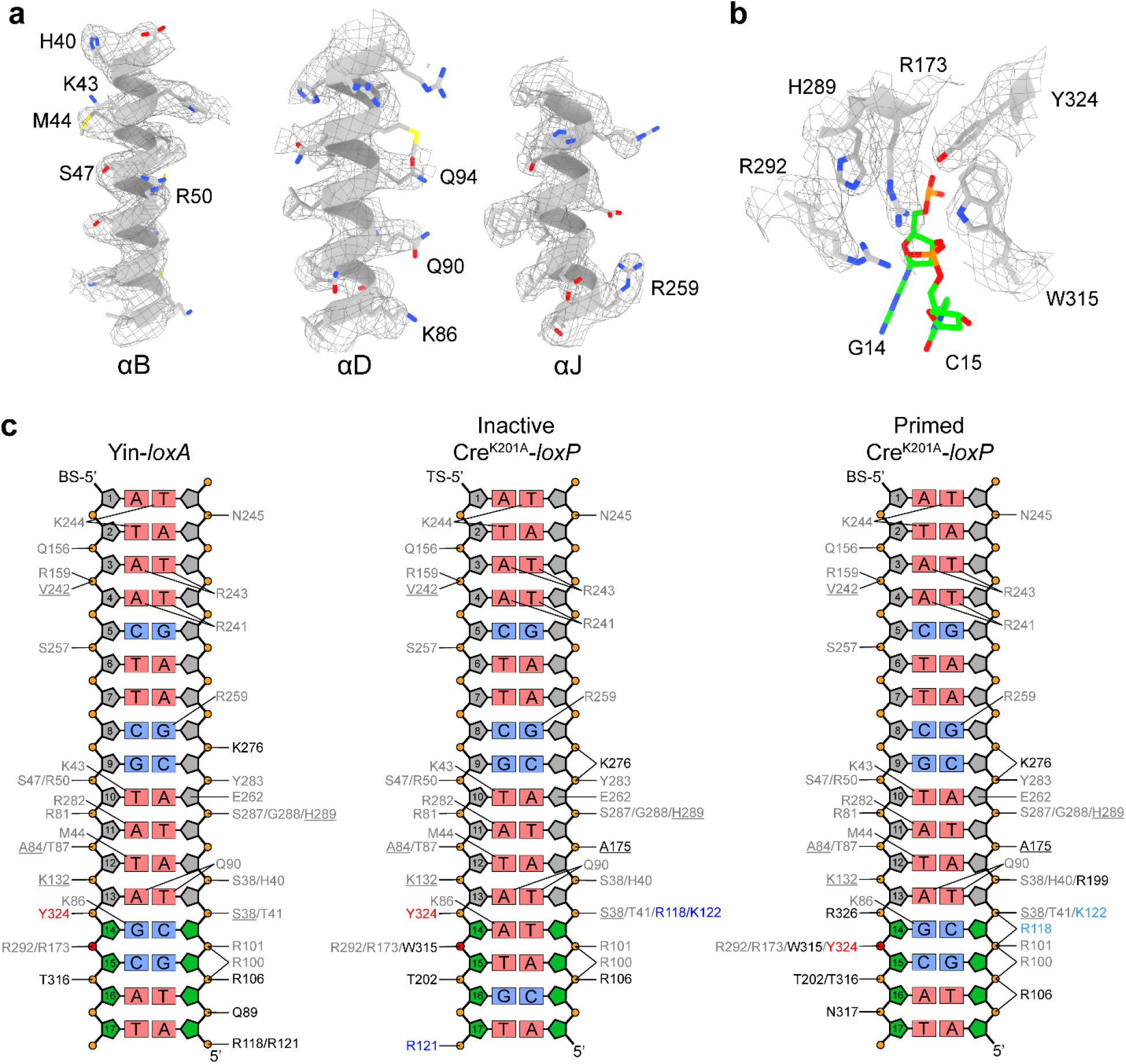
l*o*xP DNA recognition by Cre. **(a,b)** Model to map fit for helices inserted into the major grooves **(a)**, and the active site of the Cre^Yin^-loxA hairpin Cryo-EM structure **(b)**. **(c)** Protein-DNA contacts (defined as protein-DNA distances ≤ 3.5 Å) observed in the monomer and tetramer structures. Contacts colored in grey are conserved between each half-site, contacts in black vary across the half-sites, light blue contacts are protein DNA contacts across the synapse, and contacts in blue originate from the other protomer bound to the same loxP site. RBE ribose rings are colored grey and spacer ribose rings are colored green. The catalytic tyrosine is colored red and the scissile phosphate in magenta.

**Supplemental Movie 1.**
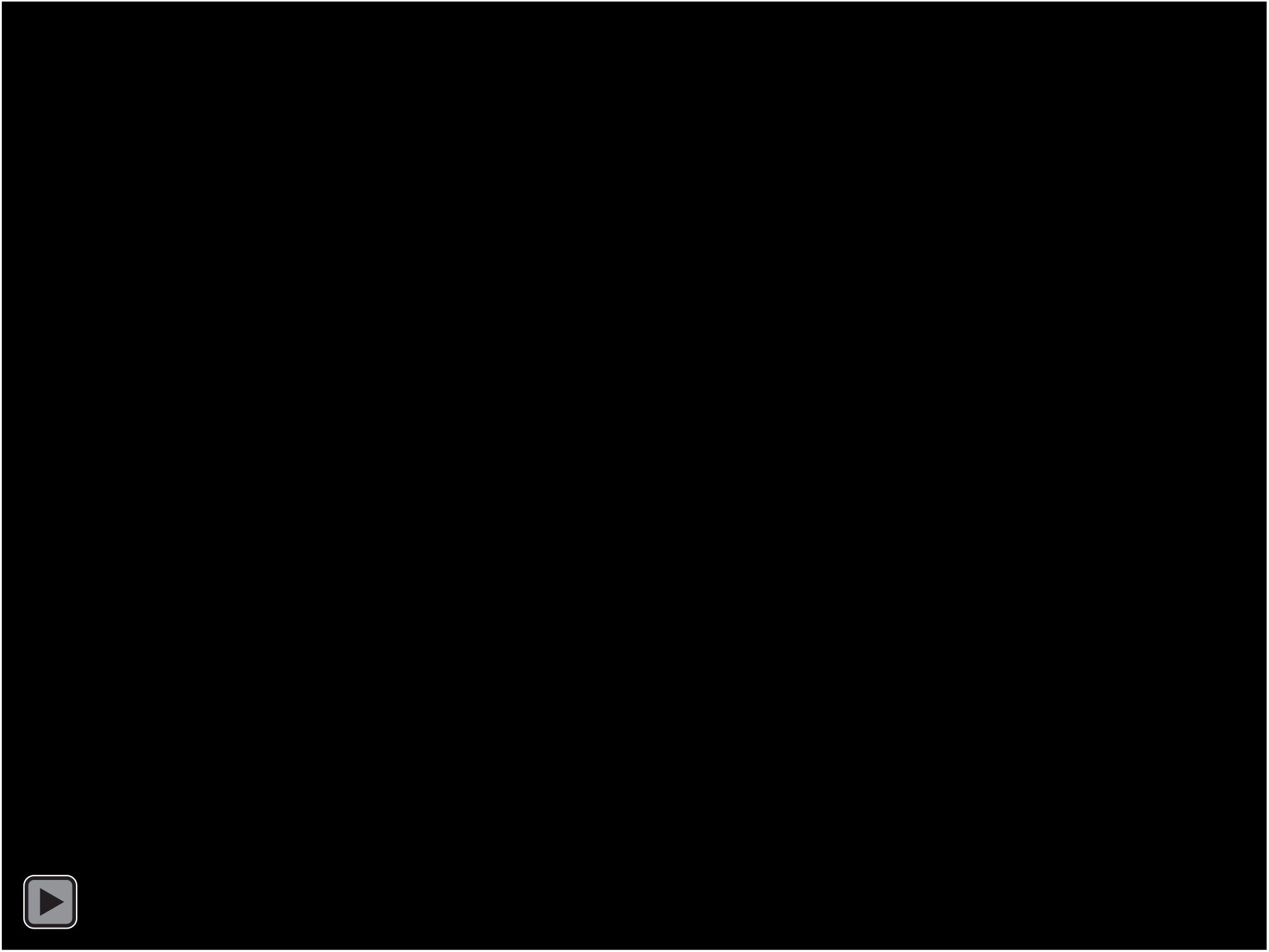
Principal component 1 from 3D variance analysis of the Cre^K201A^-*loxP* tetrameric complex cryo-EM data. This movie highlights the conformational changes associated with site-selection and activation of Cre tetrameric complexes.

**Supplemental Table 1.**
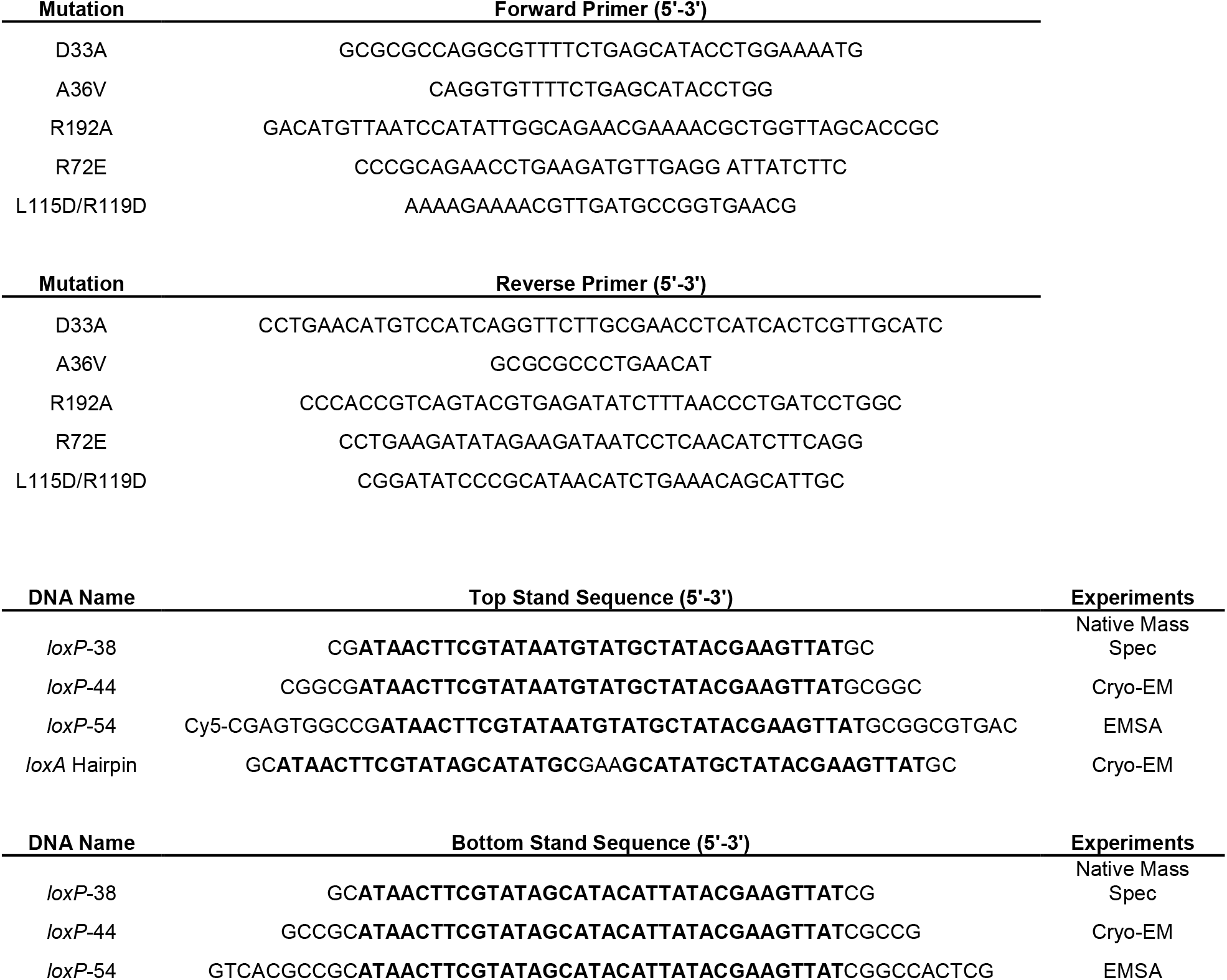
Mutagenic primers and DNA oligos. *loxP* RBE’s shown in bold.

**Supplemental Table 2.** Predicted and observed masses of individual components used in native mass spectrometry experiments. Symbols match those in Supplementary Figure 3.

**Cre<sup>Yin</sup>**
| Symbol | Mass (Da) | Intensity | Expected | Difference (Da) | Species |
| --- | --- | --- | --- | --- | --- |
| ○ | 22086 | 3 | 38307 | -16221 |  |
| ▽ | 37542 | 1 | 38307 | -765 |  |
| △ | 38307 | 100 | 38307 | 0 | 1[Y<sup>in</sup>] |
| ▷ | 38343 | 27 | 38307 | 36 |  |
| □ | 38372 | 12 | 38307 | 65 |  |
| ◇ | 38406 | 3 | 38307 | 99 |  |
| ★ | 38435 | 1 | 38307 | 128 |  |
| ○ | 38679 | 6 | 38307 | 372 |  |
| ▽ | 38712 | 22 | 38307 | 405 |  |
| △ | 38745 | 13 | 38307 | 438 |  |
| ▷ | 38778 | 8 | 38307 | 471 |  |
| □ | 38811 | 2 | 38307 | 504 |  |

**LoxP38**
| Symbol | Mass (Da) | Intensity | Expected | Difference (Da) | Species |
| --- | --- | --- | --- | --- | --- |
| ○ | 23217 | 11 | 23350 | -133 |  |
| ★ | 23350 | 64 | 23350 | 0 | 1[LoxP38] |

**<sup>15</sup>N-Cre<sup>Yang</sup>**

| Symbol | Mass (Da) | Intensity | Expected | Difference (Da) | Species |
| --- | --- | --- | --- | --- | --- |
| ○ | 24471 | 1 | 38827 | -14356 |  |
| ▽ | 24493 | 1 | 38827 | -14334 |  |
| △ | 38073 | 2 | 38827 | -754 |  |
| ▷ | 38097 | 3 | 38827 | -730 |  |
| □ | 38122 | 2 | 38827 | -705 |  |
| ◇ | 38143 | 1 | 38827 | -684 |  |
| ★ | 38769 | 1 | 38827 | -58 |  |
| ○ | 38827 | 21 | 38827 | 0 | 1[Yang<sup>15</sup>N] (~97%) |
| ▽ | 38848 | 75 | 38827 | 21 |  |
| △ | 38871 | 100 | 38827 | 44 |  |
| ▷ | 38895 | 90 | 38827 | 68 |  |
| □ | 38917 | 62 | 38827 | 90 |  |
| ◇ | 38939 | 33 | 38827 | 112 |  |
| ★ | 38961 | 10 | 38827 | 134 |  |
| ○ | 39264 | 1 | 38827 | 437 |  |
| ▽ | 39287 | 1 | 38827 | 460 |  |

**Supplemental Table 3.**
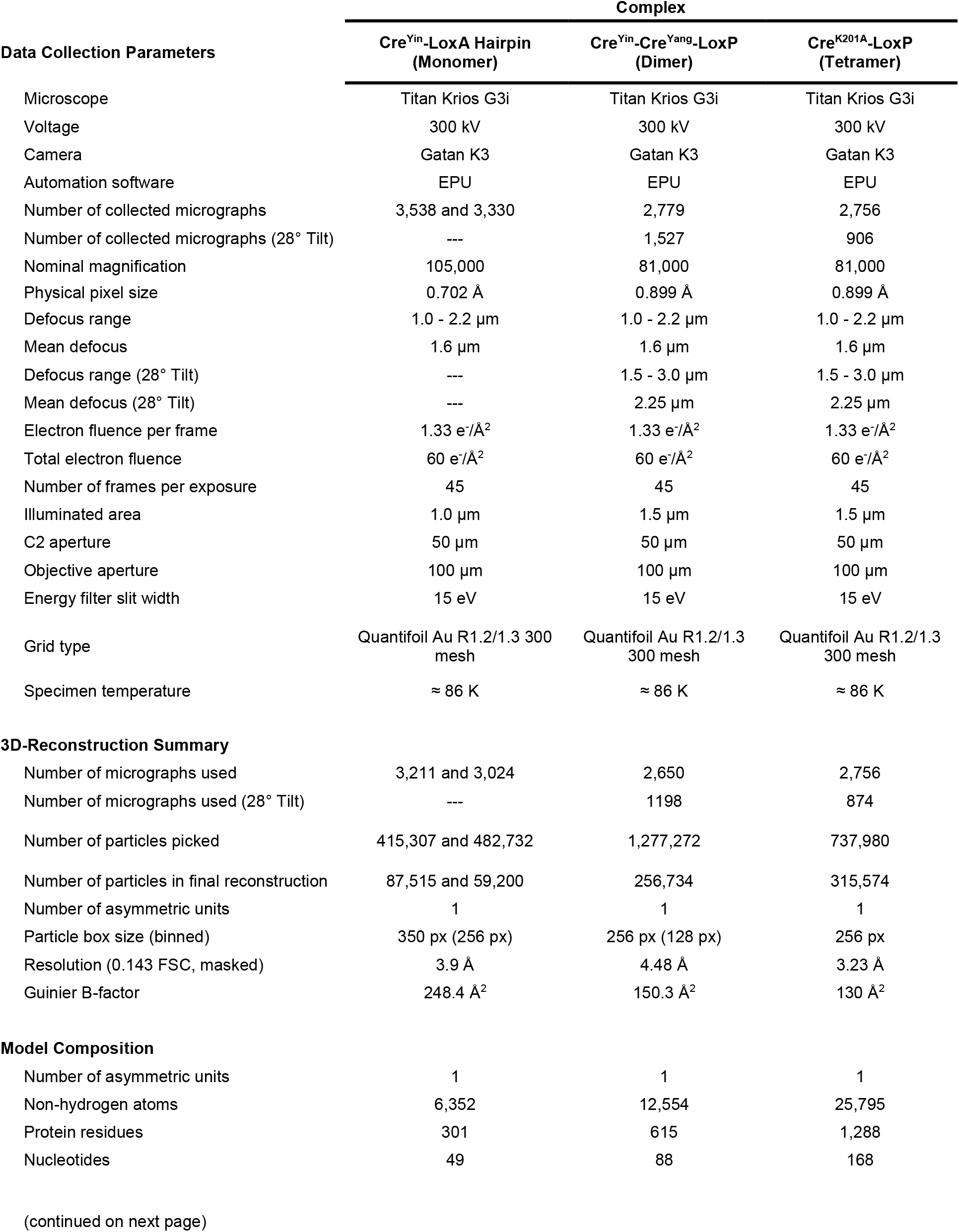
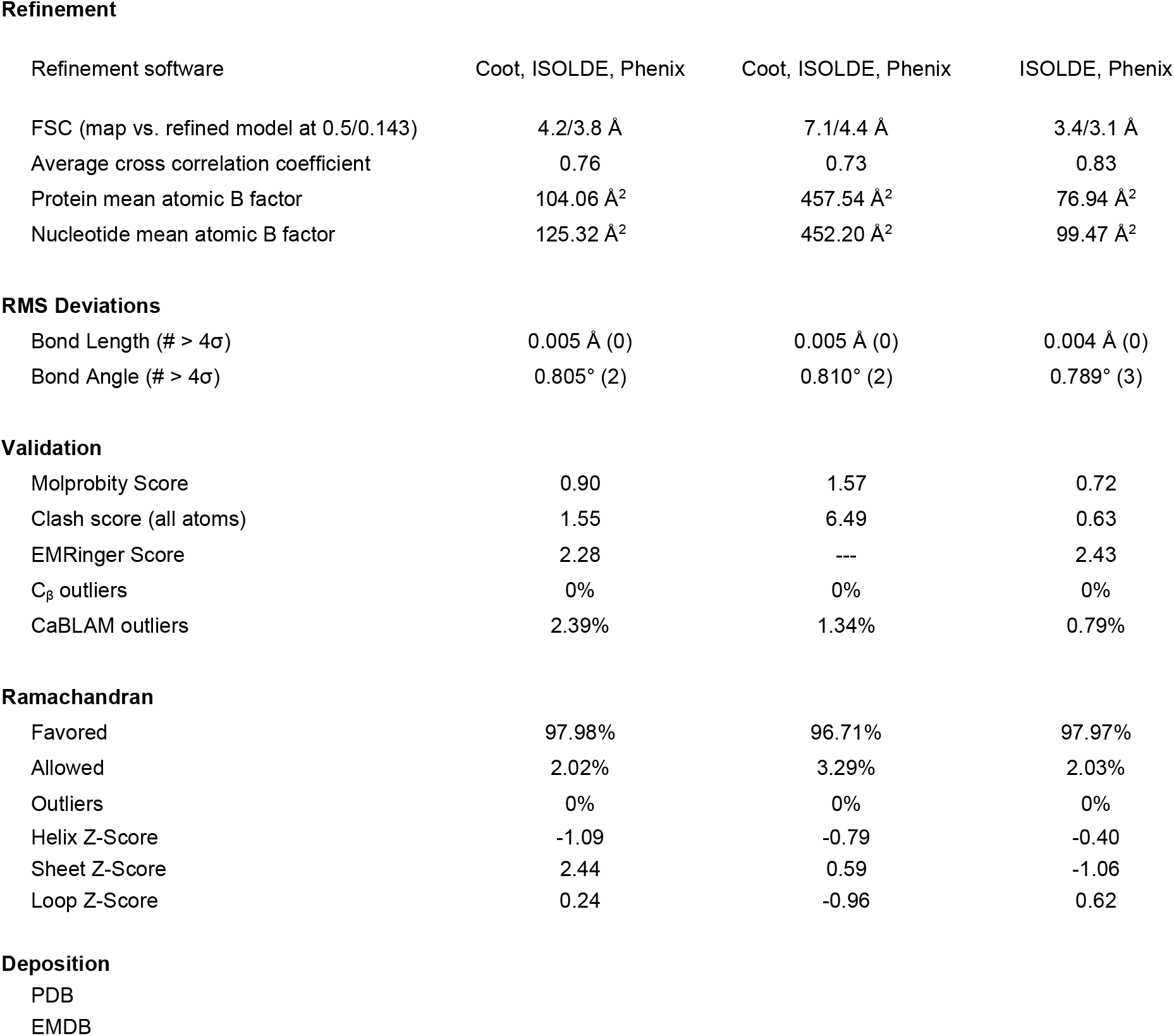
Data collection and model building summary for Cryo-EM experiments.

| Data Collection Parameters | Complex |  |  |
| --- | --- | --- | --- |
|  | Cre<sup>Yin</sup>-LoxA Hairpin<br>(Monomer) | Cre<sup>Yin</sup>-Cre<sup>Yang</sup>-LoxP<br>(Dimer) | Cre<sup>K201A</sup>-LoxP<br>(Tetramer) |
| Microscope | Titan Krios G3i | Titan Krios G3i | Titan Krios G3i |
| Voltage | 300 kV | 300 kV | 300 kV |
| Camera | Gatan K3 | Gatan K3 | Gatan K3 |
| Automation software | EPU | EPU | EPU |
| Number of collected micrographs | 3,538 and 3,330 | 2,779 | 2,756 |
| Number of collected micrographs (28° Tilt) | --- | 1,527 | 906 |
| Nominal magnification | 105,000 | 81,000 | 81,000 |
| Physical pixel size | 0.702 Å | 0.899 Å | 0.899 Å |
| Defocus range | 1.0 - 2.2 µm | 1.0 - 2.2 µm | 1.0 - 2.2 µm |
| Mean defocus | 1.6 µm | 1.6 µm | 1.6 µm |
| Defocus range (28° Tilt) | --- | 1.5 - 3.0 µm | 1.5 - 3.0 µm |
| Mean defocus (28° Tilt) | --- | 2.25 µm | 2.25 µm |
| Electron fluence per frame | 1.33 e<sup>-</sup>/Å<sup>2</sup> | 1.33 e<sup>-</sup>/Å<sup>2</sup> | 1.33 e<sup>-</sup>/Å<sup>2</sup> |
| Total electron fluence | 60 e<sup>-</sup>/Å<sup>2</sup> | 60 e<sup>-</sup>/Å<sup>2</sup> | 60 e<sup>-</sup>/Å<sup>2</sup> |
| Number of frames per exposure | 45 | 45 | 45 |
| Illuminated area | 1.0 µm | 1.5 µm | 1.5 µm |
| C2 aperture | 50 µm | 50 µm | 50 µm |
| Objective aperture | 100 µm | 100 µm | 100 µm |
| Energy filter slit width | 15 eV | 15 eV | 15 eV |
| Grid type | Quantifoil Au R1.2/1.3 300 mesh | Quantifoil Au R1.2/1.3 300 mesh | Quantifoil Au R1.2/1.3 300 mesh |
| Specimen temperature | ≈ 86 K | ≈ 86 K | ≈ 86 K |
| **3D-Reconstruction Summary** |  |  |  |
| Number of micrographs used | 3,211 and 3,024 | 2,650 | 2,756 |
| Number of micrographs used (28° Tilt) | --- | 1198 | 874 |
| Number of particles picked | 415,307 and 482,732 | 1,277,272 | 737,980 |
| Number of particles in final reconstruction | 87,515 and 59,200 | 256,734 | 315,574 |
| Number of asymmetric units | 1 | 1 | 1 |
| Particle box size (binned) | 350 px (256 px) | 256 px (128 px) | 256 px |
| Resolution (0.143 FSC, masked) | 3.9 Å | 4.48 Å | 3.23 Å |
| Guinier B-factor | 248.4 Å<sup>2</sup> | 150.3 Å<sup>2</sup> | 130 Å<sup>2</sup> |
| **Model Composition** |  |  |  |
| Number of asymmetric units | 1 | 1 | 1 |
| Non-hydrogen atoms | 6,352 | 12,554 | 25,795 |
| Protein residues | 301 | 615 | 1,288 |
| Nucleotides | 49 | 88 | 168 |

**Refinement**
| Refinement software | Coot, ISOLDE, Phenix | Coot, ISOLDE, Phenix | ISOLDE, Phenix |
| --- | --- | --- | --- |
| FSC (map vs. refined model at 0.5/0.143) | 4.2/3.8 Å | 7.1/4.4 Å | 3.4/3.1 Å |
| Average cross correlation coefficient | 0.76 | 0.73 | 0.83 |
| Protein mean atomic B factor | 104.06 Å<sup>2</sup> | 457.54 Å<sup>2</sup> | 76.94 Å<sup>2</sup> |
| Nucleotide mean atomic B factor | 125.32 Å<sup>2</sup> | 452.20 Å<sup>2</sup> | 99.47 Å<sup>2</sup> |
| **RMS Deviations** |  |  |  |
| Bond Length (# > 4σ) | 0.005 Å (0) | 0.005 Å (0) | 0.004 Å (0) |
| Bond Angle (# > 4σ) | 0.805° (2) | 0.810° (2) | 0.789° (3) |
| **Validation** |  |  |  |
| Molprobity Score | 0.90 | 1.57 | 0.72 |
| Clash score (all atoms) | 1.55 | 6.49 | 0.63 |
| EMRinger Score | 2.28 | --- | 2.43 |
| C<sub>β</sub> outliers | 0% | 0% | 0% |
| CaBLAM outliers | 2.39% | 1.34% | 0.79% |
| **Ramachandran** |  |  |  |
| Favored | 97.98% | 96.71% | 97.97% |
| Allowed | 2.02% | 3.29% | 2.03% |
| Outliers | 0% | 0% | 0% |
| Helix Z-Score | -1.09 | -0.79 | -0.40 |
| Sheet Z-Score | 2.44 | 0.59 | -1.06 |
| Loop Z-Score | 0.24 | -0.96 | 0.62 |

**Supplemental Table 4.** Buried surface area for each protein-protein and protein-DNA interface of Cre assembly intermediates.

| Interface | Buried Surface Area (Å<sup>2</sup>) |
| --- | --- |
| **Protein - DNA** |  |
| Monomer | 2,138.3 |
| Cre<sup>Yin</sup> | 2,027.9 |
| Cre<sup>Yang</sup> | 2,111.3 |
| Tetramer - Inactive | 2,156.9 |
| Tetramer - Primed | 2,530.3 |
| **Protein - Protein** |  |
| Monomer | 0 |
| Dimer | 695.4 |
| Tetramer - Synapse | 1,468.6 |
| Tetramer - Along<br>DNA | 1,842.2 |

